# Microbial Metabolic Enzymes, Pathways and Microbial Hosts for Co-Metabolic Degradation of Organic Micropollutants in Wastewater

**DOI:** 10.1101/2024.09.04.611302

**Authors:** Boyu Lyu, Bharat Manna, Xueyang Zhou, Ivanhoe K. H. Leung, Naresh Singhal

## Abstract

Organic micropollutants (OMPs) in wastewater present significant environmental challenges, but effective removal strategies are hindered by our limited understanding of their co-metabolic biodegradation. We aim to elucidate the microbial enzymes, metabolic pathways, and community members involved in OMP co-metabolic degradation, thereby paving the way for more effective wastewater treatment strategies. We integrated multi-omics (metagenomics, metaproteomics, and metabolomics) and functional group analysis to investigate 24 OMPs under three aeration conditions. Our findings reveal that oxidoreductases, particularly cytochrome P450s and peroxidases, are crucial for recalcitrant OMPs containing halogen groups (-Cl, -F) like fluoxetine and diuron. Hydrolases, including amidases, are instrumental in targeting amide-containing (-CONH₂) OMPs such as bezafibrate and carbamazepine. Regarding microbial metabolism involved in OMP co-metabolic degradation, we found that amino acid metabolism is crucial for degrading amine-containing (-NH₂) OMPs like metoprolol and citalopram. Lipid metabolism, particularly for fatty acids, contributes to the degradation of carboxylic acid (-COOH) containing OMPs such as bezafibrate and naproxen. Finally, with *Actinobacteria*, *Bacteroidetes*, and *Proteobacteria* emerging as primary contributors to these functionalities, we established connections between OMP functional groups, degradation enzymes, metabolic pathways, and microbial phyla. Our findings provide generalized insights into structure-function relationships in OMP co-metabolic degradation, offering the potential for improved wastewater treatment strategies.

**Graphical abstract:** 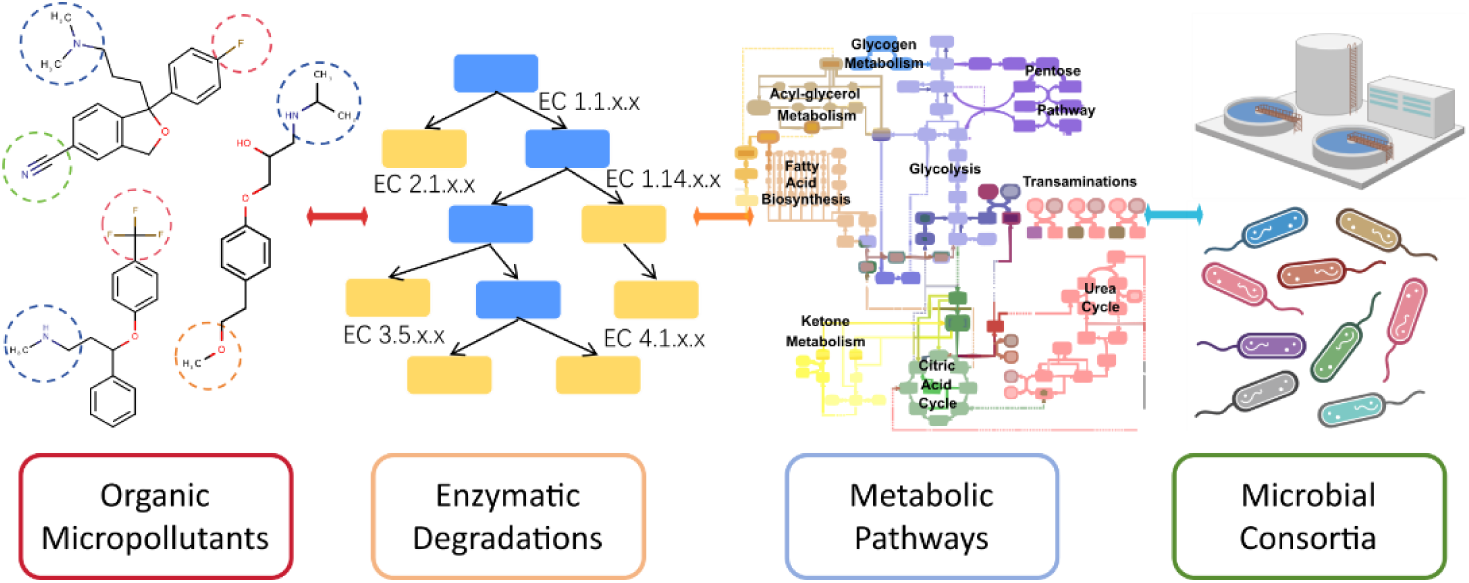

## 1. Introduction

The inadequate removal of bioactive and toxic OMPs by municipal WWTPs has led to their rampant discharge into aquatic ecosystems (Mao et al., 2022; Wang et al., 2022a). These OMPs, including analgesics, antibiotics, and steroids, pose a significant ecological threat worldwide (Wang et al., 2018). Despite their low concentrations in receiving waters (Falås et al., 2016), OMPs show environmental persistence and can inflict adverse ecotoxicological effects on exposed organisms via endocrine disruption, impaired immunity, neurological impacts, and reduced reproductive capacity (Bilal et al., 2019; Correia et al., 2023; Shao et al., 2019). The global health and economic costs associated with exposure to environmental chemicals, including OMPs, are estimated to reach a staggering ∼$6.2 trillion annually (Grandjean and Bellanger, 2017). However, despite the recognized urgency, progress in the development of effective OMP removal systems remains slow (Kulandaivelu et al., 2020; Petrie et al., 2014).

Biodegradation is commonly used to degrade chemicals in wastewater. Microorganisms can facilitate the transformation of OMPs via direct metabolic pathways or indirect co-metabolic processes (Kennes-Veiga et al., 2022a). As OMPs are typically present at trace levels in wastewater, the concentration of OMPs is unlikely to be high enough to meet the primary metabolic requirements of microorganisms for sustaining energy and growth. Hence, co-metabolic transformation is likely the dominant pathway (Kennes-Veiga et al., 2022a). In co-metabolism, microbes leverage the promiscuous nature of enzymes to catalyze the transformation of OMPs that are structurally similar to the primary substrates (Oliveira et al., 2019). So far, our understanding of the specific enzymes, metabolic pathways, and microbial communities involved in this process remains limited (Oliveira et al., 2019).

Previous studies have demonstrated that various bacteria and enzymes could degrade OMPs. For example, it has been demonstrated that bacteria such as *Methylobacter* and *Nitrosomonas* could readily degrade aromatic OMPs like caffeine and 2,4-dichlorophenoxyacetic acid (Wang et al., 2022b), while others like *Mycobacterium, Thauera,* and *Stentrophomonas* harbor various documented biodegradation genes such as cytochrome P450s that may facilitate the degradation of aromatic pollutants including atrazine and carbendazim (Fang et al., 2018; Gao et al., 2024). At the enzyme level, amidases (EC 3.5.1.-) have been shown to catalyze the hydrolysis of amide-containing OMPs (Achermann et al., 2018). Acetate kinase (EC 2.7.2.1) may facilitate the degradation of naproxen and diclofenac (Gonzalez-Gil et al., 2017). Additionally, ammonia monooxygenase (EC 1.14.99.39) possesses the ability to detoxify antibiotics like cephalexin by targeting their β-lactam function group (Wang et al., 2019). Finally, different treatment conditions have also shown to influence microbial OMP degradation capabilities by altering heterotrophic activity and microbial communities (Harb et al., 2016; Kennes-Veiga et al., 2021).

However, these assessments are often disconnected, lacking a comprehensive understanding of the OMP structure-enzyme relationships and the cellular metabolic processes in which these enzymes naturally operate. As a result, we lack an understanding of the cellular metabolisms of specific microbes that could be targeted to overproduce enzymes for faster OMP degradation. Furthermore, with over 350,000 OMPs present in waterbodies globally (Wang et al., 2020), it is necessary that we generalize the existing biodegradation knowledge to make it applicable to uncharacterized or less studied OMPs. We hypothesize that using chemical functional groups as fingerprints, we can relate OMPs to biotransformation pathways and link the transformation enzymes to cellular metabolisms and bacterial hosts. We adopt a multi-omics approach to connect the metagenome, enzyme pools, and metabolites within the WWTP microbiome (Johnson et al., 2015). Our goal is to provide comprehensive insights into the co-metabolic degradation of OMPs and pave the way for developing more targeted and effective wastewater treatment strategies.

## 2. Materials and Methods

### 2.1 Reactor Operation and Parameters

The activated sludge used in this study was sourced from the Māngere Wastewater Treatment Plant in Auckland, New Zealand. The experiment consisted of six identical 1L bioreactors operated simultaneously at a room temperature of 20 ± 1°C. Each bioreactor was subjected to different aeration conditions over a 48-hour incubation period (Figure S1). In condition A, the dissolved oxygen (DO) concentration was maintained at a constant level of 2.0 ±0.2 mg/L. In condition B, DO varied in 3-minute cycles, where the DO increased from 0 to 2.0 ±0.2 mg/L and then decreased to 0 mg/L within each cycle. In condition C, DO underwent cyclic changes in 6-minute cycles. In each cycle, the DO increased from 0 to 2.0 ± 0.2 mg/L and then decreased to 0 mg/L within 3 minutes, followed by a 3-minute period where the DO was maintained at 0 mg/L. Conditions D, E, and F followed similar operation strategies as conditions A, B, and C, respectively, but with a different DO range of 0∼8 ± 0.2mg/L. Rapid DO increases were achieved in the three conditions that employed a fixed air flow rate of 0.5 L/min with a mass flow controller and compressed air. Initially, the reactors contained 2.75 g/L of mixed liquor suspended solids (MLSS) activated sludge, 3.84 g/L of NaHCO3 as an inorganic carbon source, and 1 mL of a trace element solution (Table S1). 50ml Synthetic wastewater (Table S1) was continuously fed into the reactors using a syringe pump and thoroughly mixed using a magnetic stirrer.

### 2.2 Organic Micropollutants Sampling and Analysis

OMPs were purchased from (AK Scientific, USA) with the highest analytical purity. The working solution was prepared prior to the experiment, and the 1L reactor was spiked with 10μg/L of the 24 OMPs. To extract the remaining OMPs, solid phase extraction (SPE) was performed using OASIS Prime HLB cartridges (Waters, Milford, MA, U.S.A.) with a standard method (Česen et al., 2018). Briefly, at the end of the cultivation period, 200 mL of the bioreactor culture was collected and centrifuged (10,000g, 4 °C, 20 min) to remove suspended particles and cell pellets. The supernatant was acidified to pH=2 using 0.1M HCl and then passed through cartridges, washed with 5% methanol, and finally, the OMPs were eluted with 100% methanol. The extracted OMPs were analyzed using a Shimadzu 8040 LC-MS/MS instrument (Shimadzu, Japan) with a C18 column (2.1 × 100 mm2, particle size 1.8 μm, Agilent Technologies, Germany). A binary gradient of 0.1% formic acid in deionized water (mobile phase A) and 0.1% formic acid in methanol (mobile phase B) were used for both ESI + and ESI - model. The flow rate for both modes was 0.25 mL/min for 20 mins, and internal standards were used to minimize matrix effects (Table S2). Limits of detection and quantification were set with signal-to-noise ratios of 10. OMP removal efficiency was calculated for 48 hours against autoclaved sludge to minimize abiotic effects with statistical analysis via Prism 9, respectively.

### 2.3 DNA Extraction and Metagenomics Analysis

DNA was extracted using DNeasy PowerSoil Pro Kit (QIAGEN, Germany). Metagenomics processing, including DNA quality assessment and library preparation, was performed at Auckland Genomic Center (Auckland, NZ). Sequencing utilized the HiSeqX platform with 2×150 bp paired-end shotgun sequencing. Preprocessing employed Trimmomatic (Bolger et al., 2014) with specific quality thresholds. SqueezeMeta v1.5.2 (Tamames and Puente-Sánchez, 2019) pipeline was used for metagenomic taxonomic and functional profiling, including co-assembly with Megahit. RNAs and ORFs were predicted using Barrnap and Prodigal, respectively. Taxonomic assignments used NCBI GenBank nr database (Clark et al., 2016) with rank-specific identity thresholds. Functional assignments utilized the KEGG database with Diamond (Buchfink et al., 2014). Detailed methods, including specific software versions, parameter settings, and preprocessing steps, are provided in Supplementary Text 1. Raw metagenomes are available at European Nucleotide Archive with data identifier PRJEB74089.

### 2.4 Metaproteomic analysis

Metaproteomic analysis was performed on 5 mL sludge samples under various experimental conditions, with three biological replicates per condition. Sludge samples were pelleted, washed, and protein extraction was conducted by sonication, and centrifugation. Purification was achieved with SpeedBead carboxylate-modified E3 and E7 magnetic particles (Sera-Mag, USA) to yield 150μg protein for each sample with EZQ protein assay kit to quantify the protein concentration. The protein extract was then reduced with 5 mM dithiothreitol (DTT), alkylated with 15 mM iodoacetamide, and digested with 1.5 µg trypsin. Peptides were cleaned by solid-phase extraction (Oasis Prime HLB 1cc, 30 mg), and concentrated using a speed vacuum. Peptide samples (10 µL) were analyzed by nano LC-MS/MS using a NanoLC 400 UPLC system and a TripleTOF 6600 Quadrupole-Time-of-Flight mass spectrometer. Data were searched against a metagenomics-derived database using MetaProteomeAnalyzer and X-tandem, with a 1% false discovery rate filter. The resulting group file was uploaded to MetaProteomeAnalyzer for metaproteomics analysis and clustering. Enzyme regulation information was obtained from RegulonDB v12.0. Raw metaproteomics data is available at ProteomeXchange with dataset identifier PXD044490.

### 2.5 Metabolomics analysis

The relative abundance of intracellular metabolites in activated sludge was determined using gas chromatography/mass spectrometry (GC/MS), with three biological and two technical replicates analyzed for each experimental condition. Metabolites were extracted by adding 1:1 methanol-water solution and an internal standard (2,3,3,3-d4-alanine) to the sludge pellets, followed by three cycles of freeze-thawing, vigorous shaking, and centrifugation. The metabolite extract was reconstituted in sodium hydroxide solution, supplemented with pyridine and methanol, and derivatized with methyl chloroformate. The derivatized sample was then analyzed using a gas chromatography–mass spectrometry (GC-MS) system equipped with a ZB-1701 GC capillary column. GC/MS chromatograms were deconvoluted by AMDIS software, and mass spectra were compared to an in-house MS library for metabolite identification. Data filtering was performed using the MassOmics R package and metabolite abundance values were normalized with the internal standard and baseline calibrated. Raw metabolomics is available at MetaboLights with the dataset identifier MTBLS8331.

### 2.6 Biotransformation Rules Extraction and Analysis

EAWAG (EAWAG-BBD, 2024) and EnviPath (Wicker et al., 2016) databases were used to predict biotransformation pathways for OMPs, including all aerobic likelihoods, and genes with EC numbers at sub-subclasses level for each biotransformation rule were extracted and summarized. For each OMP, the related gene candidate was extracted from metagenomics data using RStudio33 (R version 4.3.2) and Excel Visual Basic for Applications (VBA) code. In brief, the EC numbers at the third digit were summarized from EnviPath, and genes with the same ID identified from metagenomic data were extracted into new Excel sheets via the search and copy function. The general data process frame is shown in Figure S2. A list of relevant biotransformation rules (btrules), relevant functional groups, and types of reactions is provided in Table S3; for all 24 OMPs the corresponding EC numbers are provided in Table S3. Heatmaps are generated by the R package of ggplot2.

### 2.7 Statistical Analysis of Gene Accumulation, OMP Degradation, and Microbial Consortium

Pearson correlation coefficients (Pearson’s r) were calculated using R (version: 3.6.3) between degradation of OMPs and gene abundance (TPM) across three test conditions of A, B, and C. Genes with high correlation coefficients (r > 0.6) were considered for potential OMP biotransformation. In the EC number classification scheme, enzymatic reaction types are typically defined at the third level (sub-subclasses) of the four-digit EC numbers, whereas the fourth digit characterizes substrate specificity. We performed correlation analysis at the level of individual ECs (fourth-level EC numbers). Using OmicStudio (Lyu et al., 2023), correlation coefficients and hierarchical clustering were performed to construct a correlation heatmap using Euclidean distances and complete linkages. Mantel tests were utilized to identify the relationship between microbial abundance at the phyla level and gene accumulations under specific EC sub subclasses with statistical significance of P< 0.05 and Rho > 0.5. The visual linkage was achieved via SankeyMATIC.

## 3. Results and Discussion

### 3.1 Removal of Organic Micropollutants

Our analysis of 24 OMPs spiked in sludge-seeded reactors revealed varying degrees of degradation across all conditions (Figure 1). Eight compounds were nearly completely removed (>95% removal), five exhibited >70% removal, six showed removal between 70% and 40%, and five demonstrated <40% degradation. Structural analyses (Table S3) indicated that electron-donating functional groups like amide and hydroxyl could enhance OMP degradation, exemplified by acetaminophen (>95% degradation). Our work found that the electronegative functional groups (such as -Cl and -F) appeared to impede biodegradation. For example, diuron and fluoxetine, which possess halogen groups (-Cl or -F), achieved <40% degradation. These findings align with previous studies (Rich et al., 2022;Falås et al., 2016) that reported higher biodegradability for OMPs like acetaminophen, bezafibrate, and clarithromycin, while diuron, fluoxetine, and venlafaxine are more resistant to degradation.

**Figure 1.**
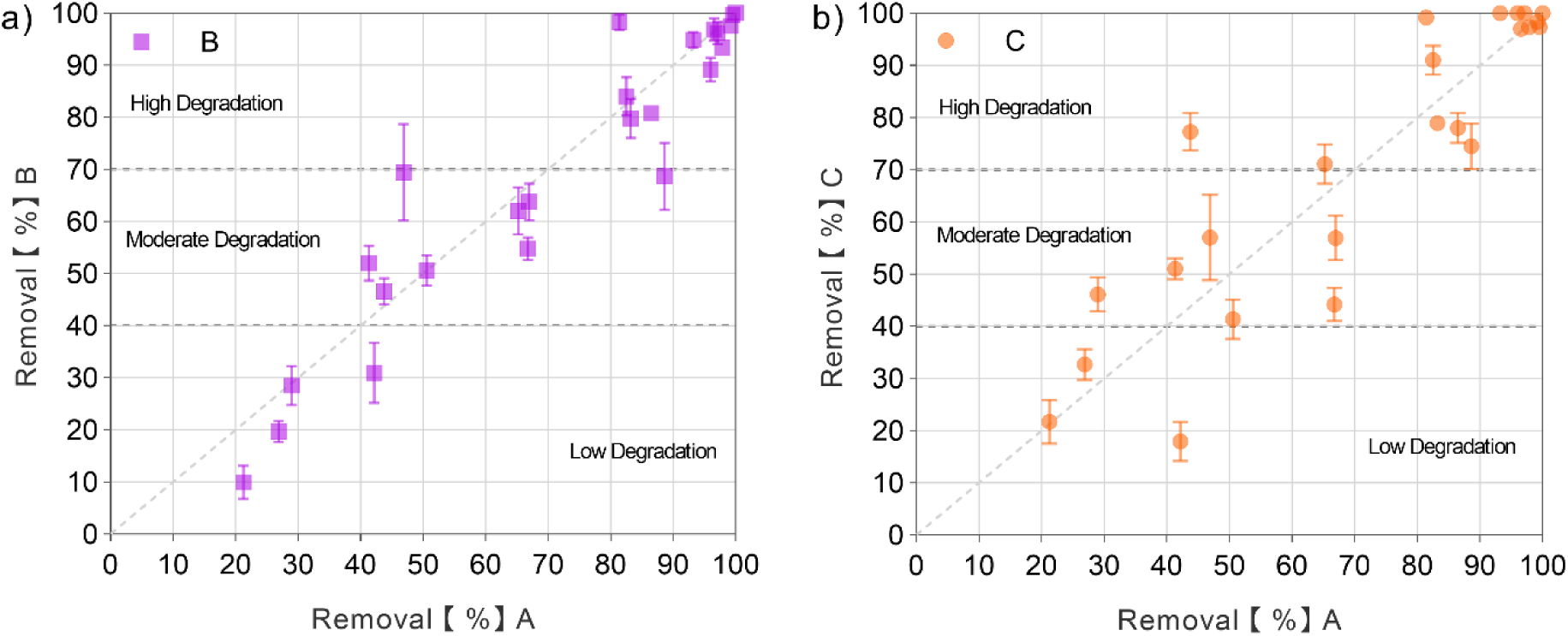
Comparison of OMP removal under different treatment conditions. (a) Removal efficiency of 24 OMPs under conditions A versus B. (b) Removal efficiency of 24 OMPs under conditions A versus C.

To further explore OMP biodegradation, we investigated three different conditions (A, B and C, see methods). Given the low concentration of OMPs (10 μg/L) in our experiments, co-metabolic degradation likely served as the primary mechanism for OMP removal (Kennes-Veiga et al., 2022a). Condition B, compared to condition A, resulted in greater degradation for gemfibrozil (26 ± 6%), clarithromycin (18 ± 2%), and venlafaxine (12 ± 3%). Notably, these three contaminants share an ether linkage (-O-) in their chemical structure (Table S3). Previous studies have documented enzymatic degradation through this functional group, such as venlafaxine degradation via cytochrome P450s (Magalhães et al., 2014), and erythromycin (precursor of clarithromycin) degradation by microbial glycoside hydrolases (Ren et al., 2021). In contrast, condition C showed increased degradation for metoprolol (35 ± 4%), diuron (21 ± 4%), clarithromycin (19 ± 1%), venlafaxine (11 ± 2%), fluoxetine (9 ± 3%), and citalopram (9 ± 3%) (One-way ANOVA, P<0.05, Figure 1). These OMPs are characterized by the presence of amine groups (-NH2, -NH-, -N-). Biodegradation through this functional group has been previously observed in diuron (Singh and Singla, 2019), and venlafaxine (Llorca et al., 2019). Various P450s have shown the capability to facilitate N-dealkylation from the amine group in fluoxetine (Deodhar et al., 2021). Multiple functional groups in each OMP suggest that their degradation involves complex networks of microbial enzymes, metabolic pathways, and microbial communities working together (Bains et al., 2019; Lim et al., 2020; Rios-Miguel et al., 2023). To systematically unravel the governing factors for OMP degradation, the functional groups on each OMP shall play a crucial role in understanding their unique degradation behaviors.

### 3.2 Structural features and biotransformation rules mediating OMP biodegradation

To elucidate the relationship between OMP structure and biodegradation potential, we conducted a comprehensive analysis of the structural features of the 24 OMPs and their potential biotransformation pathways. Our analysis revealed that the most prevalent structural features include aromatic rings, hydroxyl groups (-OH), amines (-NH2), carboxylic acids (-COOH), and amides (-CONH₂). Less common features comprise chloro (-Cl) and sulfonamide (-SO_2_NH-). To identify potential biotransformation rules (btrules), we employed the Eawag-PPD database (Table S3) (EAWAG-BBD, 2024). These literature-based btrules summarize functional groups susceptible to biodegradation and the corresponding enzyme categories (based on the enzyme classification system with enzyme commission numbers) that facilitate these transformations. For instance, amine-containing compounds can undergo oxidative removal of an R group via bt0063 (amine dealkylations), catalyzed by enzymes such as monooxygenase (EC:1.13.12.-), dehydrogenase (EC: 1.4.1.-), and aminotransferase (EC:2.6.1.-) (EAWAG-BBD, 2024). Similarly, biodegradation of amide-containing OMPs typically involves hydrolysis of the amide bond (bt0067) by enzymes like carboxylesterases (EC: 3.1.1.-), peptidases (EC: 3.4.13.-), or linear amidases (EC:3.5.1.-).

Our findings demonstrate that specific structural features significantly influence OMP degradation. For example, OMPs of benzotriazole with structural features associated with bt0005 (dihydroxylation) and bezafibrate with bt0241 (monohydroxylation) exhibited high degradation rates (>80% in condition A). Conversely, diuron, which features bt0243 (N-dealkylation), showed poor degradation (<30% in condition A). These observations align with previous research indicating that certain functional groups can either accelerate or hinder biodegradation (Rich et al., 2022). Traditionally, studies have attempted to decipher the causal genes involved in OMP degradation by identifying a limited number of transformation products (TPs) (Fenner et al., 2021). However, multiple TPs can arise simultaneously for a single OMP (Rubirola et al., 2014), and the detection method may be limited to identifying TPs if the exact mass is less than 100 Da, or the efficiency of ionizations, and if the compound is not retained on the column (Rich et al., 2022). Given these limitations, our approach considers all possible functional group-based reactions in OMP degradation and their associated enzyme categories to provide a more comprehensive view in the following sections to decipher the co-metabolic OMP degradation at trace levels.

### 3.3 Identifying metabolic Enzymes (genes) in the mixed-microbial community

Metagenomic and metaproteomics analyses were conducted to comprehensively identify the metabolic enzymes (and their corresponding genes) in the studied microbial communities across different conditions. From the metagenome, we identified 9,861 unique KEGG ortholog IDs (KO IDs), each representing the functional characteristics of the detected genes. Out of all the KO IDs, 5,612 genes were associated with different ECs that best describe the enzymatic function of the genes (Table S4). These enzymes belonged to Oxidoreductases (EC 1, n=1356), Transferases (EC 2, n=1795), Hydrolases (EC 3, n=1287), Lyases (EC:4, n=431), Isomerases (EC5, n=320), Ligases (EC6, n=234), and Translocases (EC7, n=189). To verify the gene expression, the metaproteomic analysis identified 689 unique enzyme candidates in our system (Figure S3). A total of 200 unique oxidoreductases (EC:1) have been confirmed with expression, followed by 171 transferases (EC:2), 90 hydrolases (EC: 3), 74 lyases (EC:4), and 49 isomerases (EC:5). The detection of proteins in wastewater matrix is a challenging task as the complex nature of the sample, and the presence of humic acids sharing identical chemical properties as peptides further interfered with the peptide extraction for maximum identification (Qian and Hettich, n.d.; Zhang et al., 2019). While only a portion of the total enzyme pool can be revealed, future studies could focus on optimizing the existing detection methods to capture a more complete metaproteome. It is worth noting that metagenomics captures the accumulated changes of the microbiome’s metabolic potentials and metaproteomics analysis to reveal the presence of the target enzyme, which still provides more comprehensive views of the sludge system in wastewater.

To link enzyme functions to the structural features of OMPs, we focused on the most frequent enzyme (gene) categories observed across all three conditions (n=9), including methyltransferases (268 KO IDs, EC:1.1.1.-), linear amides hydrolases (111 KO IDs, EC: 3.5.1.-), and monooxygenase (84 KO IDs, EC 1.14.13.-), which are implicated in critical reactions such as bt0063 (amine dealkylations), and monohydroxylation reactions (bt0036, bt0241, and bt0242). Using EnviPath database (Wicker et al., 2016), we identified a total of 67 unique EC sub-subclasses potentially involved in the degradation of the 24 targeted OMPs (Table S4). For each btrules, detailed categorization of the EC category with relevant KO IDs has been outlined in Table S5, with bt0063 processing 327 KO IDs, bt0023 with 376 KO IDs, bt0067 with 312 KO IDs, and bt0242 with 192 KO IDs. We then attribute the enzyme (gene) candidate to each OMP based on the available structure features. For amine-containing compounds, 1008 KO IDs were assigned for metoprolol, 580 KO IDs for fluoxetine, and 344 KO IDs for citalopram. The variation in the number of potential degrading enzymes for each OMP, due to the presence of multiple functional groups, underscores the complexity of potential OMP degradation processes. This highlights the need to further investigate the collective enzyme pools to probe key enzyme candidates for OMPs.

### 3.4 Assessing potential metabolic enzyme (genes) sub-subclasses for co-metabolic OMP degradation

Furthermore, we analyzed the correlation between the abundance of specific enzyme sub-subclasses (genes) and their btrule-related OMP degradation. This analysis was based on the assumption that higher gene abundance could be associated with higher OMP removal (Fang et al., 2018; Gao et al., 2024; Jeffries et al., 2018). Focusing on prevalent EC subclasses (EC:1.1.1.-, EC:1.14.13-, and EC:3.5.1.-). Fig 2a illustrates the Pearson’s r value (-1 to 1) between the gene abundance of all KO IDs under EC subclasses of EC:1.1.1.- and the removal of OMPs with structure features like (-OH), which are potential targets for co-metabolic degradation by EC:1.1.1.-enzymes. Interestingly, even though these OMPs are all related to dehydrogenases (EC:1.1.1.-), the enzymes within this EC subclass showed differential correlations to individual OMPs. This observation suggests OMPs having similar structure features with different natural substrates of bacterial metabolic enzymes can undergo co-metabolic transformations (Fischer and Majewsky, 2014; Gonzalez-Gil et al., 2017; Kennes-Veiga et al., 2022a).

**Figure 2.**
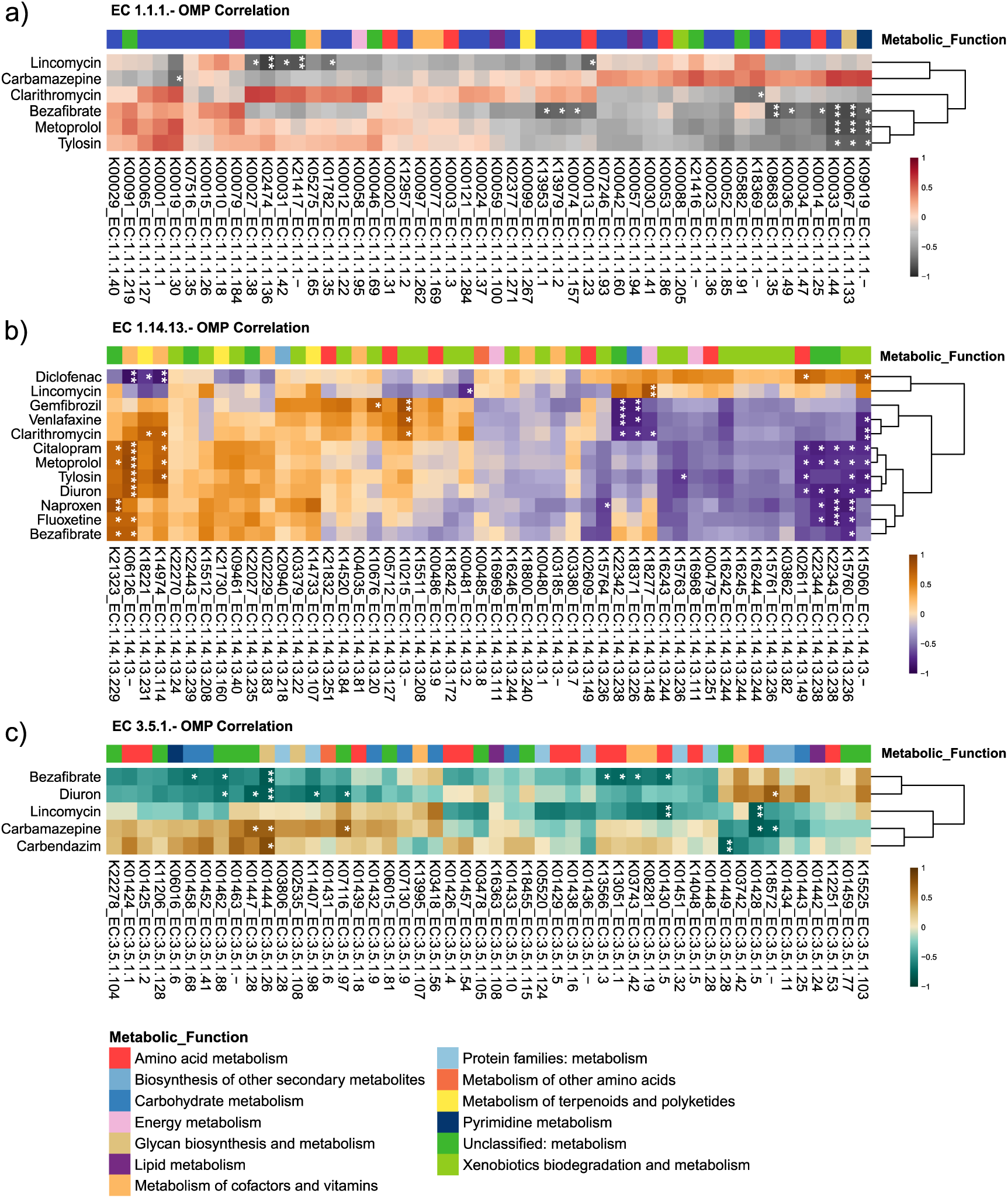
Correlation heatmaps of enzyme gene abundance and OMP degradation. (a) Correlation between EC 1.1.1.-genes (Top 50) and degradation of 6 corresponding OMPs. (b) Correlation between EC 1.14.13.-genes (Top 50) and degradation of 12 corresponding OMPs. (c) Correlation between EC 3.5.1.-genes (Top 50) and degradation of 5 corresponding OMPs. Heatmaps display Pearson’s correlation coefficients (r) ranging from -1 to 1. Row clusters indicate the natural metabolic roles of the enzymes. Asterisks (*) denote statistically significant correlations (P<0.05)

For instance, isocitrate dehydrogenase (K00031_EC:1.1.1.42) exhibits a positive correlation with clarithromycin, metoprolol, and tylosin, while showing a significant negative correlation (P<0.05) with lincomycin (Figure 2a). This indicates that isocitrate dehydrogenase, part of carbohydrate metabolism (Zhao et al., 2021) could participate in the co-metabolic degradation of positively correlated OMPs. Similarly, trimethylamine monooxygenase (K18277_EC:1.14.13.148) shows significant positive correlation (P<0.05) with lincomycin while moderately or negatively correlating with the degradation of other OMPs (Figure 2b). This enzyme, originating from energy metabolism, may contribute to the observed degradation of lincomycin. Moreover, the expression of these two enzymes was also confirmed by our metaproteomics analysis (Figure S3). The complete list of correlation results is provided in Supplementary Data1. These findings suggest that OMPs sharing the same structural feature (btrules) can potentially be degraded by different enzymes within the same EC sub-subclasses, and these enzymes originate from diverse metabolic pathways.

Intriguingly, we observed distinct degradation patterns among OMPs sharing the same functional groups in their structure. This suggests that the chemical skeleton surrounding the functional groups may also contribute to differences in degradation. For example, among the five tested OMPs containing amide groups (bezafibrate, diuron, lincomycin, carbamazepine, and carbendazim) (Figure 2c), we observed varying degrees of degradation. Bezafibrate and diuron, both with nitrogen bonded to a phenyl group, showed 93 ±3% and 46 ±4% removal, respectively. Lincomycin, which achieved 85 ± 5% degradation, has nitrogen bound to a methyl (-CH3) group with complex side chains, including a pyrrolidine ring. Carbamazepine, with 65 ±3% degradation, is unique in having nitrogen bound to two phenyl groups. Lastly, carbendazim features an amide-bound part of a benzimidazole group. These structural differences could correspond to the natural substrates of various linear amide hydrolases, potentially resulting in distinct degradation outcomes for the five OMPs. Moreover, only a subset of the amide hydrolases positively correlates with the OMPs and are potential OMP degraders. Our analysis highlights the complex relationship between enzyme abundance, OMP structure, and degradation efficiency. Future experiments should prioritize the enzyme candidate as our metaproteomics data in Figure S3 for *in vitro* examination.

### 3.5 Identifying OMP-specific degradation enzyme candidates

To refine our understanding of potential enzymes involved in individual OMP degradation, we focused on enzymes (genes) with strong Pearson correlation coefficients (r > 0.6) across all three biological triplicates. It’s important to note that for 10 OMPs showing no statistically significant differences in removal efficiency across the tested conditions, we could not proceed with correlation analysis to further refine the list of putative degrader enzymes. For metoprolol, all genes with strong correlations (r > 0.6) are summarized in Figure 3a. With the normalized gene abundances (z-scores - 2 to 2), and the natural metabolic roles for each gene classified by KEGG level 2 functions. We observed a total of 73 oxidoreductases (EC 1), 28 transferases (EC 2), 1 hydrolase (EC 3), and 8 lyases (EC4) with a strong correlation to metoprolol degradation. Metaproteomics analysis also confirms the expression of various microbial enzymes for metoprolol degradation. For instance, oxidoreductases like flavin-dependent monooxygenase (K16047, EC:1.14.14.12), which participate in xenobiotics biodegradation (Dresen et al., 2010), and transaminase (K03430, EC:2.6.1.37), both potentially target different functional groups (amine and ether) present in metoprolol. Similarly, genes strongly correlated with fluoxetine (Figure 3b) and citalopram (Figure 3c) were summarized into separate heatmaps. Fluoxetine showed correlations with 22 oxidoreductases (EC 1), 23 transferases (EC 2), 1 hydrolase (EC 3), and 4 lyases (EC4), while citalopram correlated with 30 oxidoreductases (EC 1), 11 transferases (EC 2), 1 hydrolase (EC 3), and 8 lyases (EC4). Notably, a more diverse group of oxidoreductases appears to be involved in metoprolol degradation.

**Figure 3.**
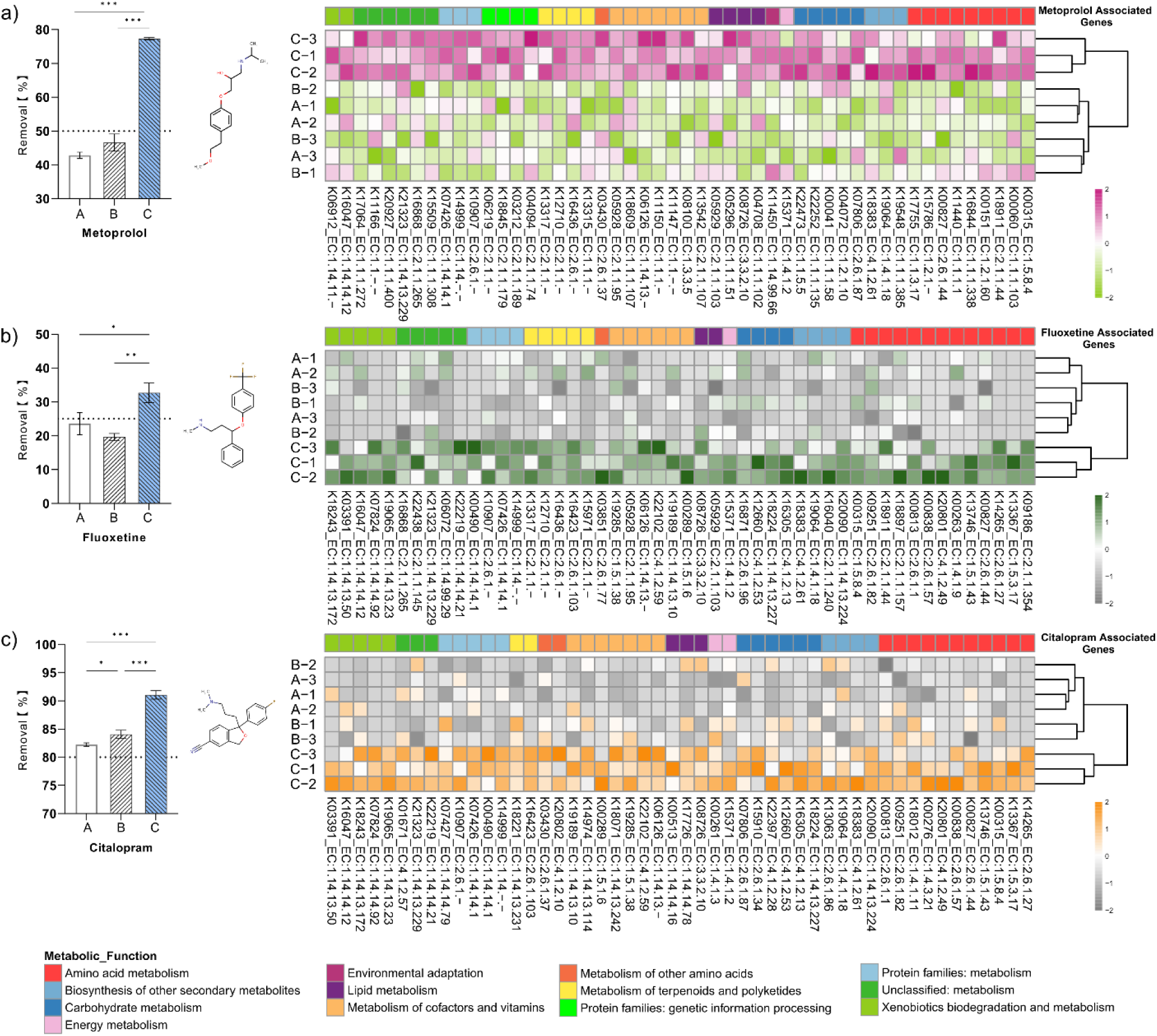
OMP degradation and associated gene candidates in abundance heatmaps (z score -2 to 2). (a) Metoprolol, (b) fluoxetine, and (c) citalopram degradation percentages after 48 hours under three aeration conditions (A, B, C). Asterisks (*) indicate significant differences (P<0.05, one-way ANOVA)

An intriguing revelation is the increased abundance of genes encoding unspecific monooxygenases and various cytochrome P450 enzymes, which correlate with the degradation of multiple OMPs (Figure 3). Unspecific monooxygenase (EC 1.14.14.1), known for its broad substrate specificity (Kennes-Veiga et al., 2022a) and capability of remediate containments in wastewater systems (Tarbajova et al., 2023), has been identified in condition C (Figure 3a) with higher gene abundance over the other two conditions. Other P450 enzymes, such as cytochrome P450 (K00490 EC:1.14.14.1), Cytochrome P450 CYP21A steroid 21-monooxygenase (K00513 EC:1.14.14.16), and cytochrome P450 family 4 (K07426 EC:1.14.14.1), with the ability to oxidize and detoxify pharmaceuticals, were also present in high abundance in condition C (Figure 3b). We suggest that the presence of different types of P450s in the system could promote denitrations and reductive dehalogenations of various OMPs (Behrendorff, 2021). Additionally, these bacterial P450s play critical roles in terpenoids, steroids, and fatty acids metabolisms (Parvez et al., 2016), pointing towards their possible co-metabolic connections with the degradation of specific OMPs (EH-Haj, 2021; Gulde et al., 2016). A total of 12 monooxygenases and dioxygenases (EC:1.14.-.-) have been confirmed with detectable levels of expression (Figure S3), and future studies could explore the catalytic mechanism in detail.

Recognizing that gene abundance accumulation in individual experiments may not provide a complete picture (Achermann et al., 2020), we conducted an independent experiment using different DO conditions (D, E, and F) with initial sludge from the same WWTP to validate our findings. We focused on degrading the same OMPs to validate the previously identified gene candidates. As shown in Figure S4, for fluoxetine, 64% of the gene candidates (27 out of 42) maintained positive correlations with degradation. For example, K16305 (EC:4.1.2.13) showed consistent positive correlations (r=0.81 for A, B, C; r=0.82 for D, E, F), as did K14999 (EC:1.14.-.-) (r=0.88 for A, B, C; r=0.65 for D, E, F). However, some genes, like K15971 (EC:2.1.1.), showed opposite correlations between experiments (r=0.87 for A, B, C; r=-0.90 for D, E, F). For other OMPs, metoprolol and citalopram showed consistent correlations for 57% and 53% of gene candidates, respectively. Future studies should evaluate seasonal effects and other treatment parameters on metabolic changes within the microbiome across different WWTPs to generalize and verify these findings.

### 3.6 Key metabolic functions governing OMP Degradation

Understanding the complex metabolic processes governing the degradation of OMPs is crucial for developing effective wastewater treatment strategies. Our analysis, based on the natural metabolic roles of positively correlated genes (enzymes) for individual OMPs, revealed that OMP removal was primarily associated with 11 key metabolic pathways, including amino acid metabolism, carbohydrate metabolism, and xenobiotics metabolism (Figure 4, Table S6). We further examined the roles of these metabolic functions in individual OMP degradation with metabolomics analysis (Figure S5).

**Figure 4.**
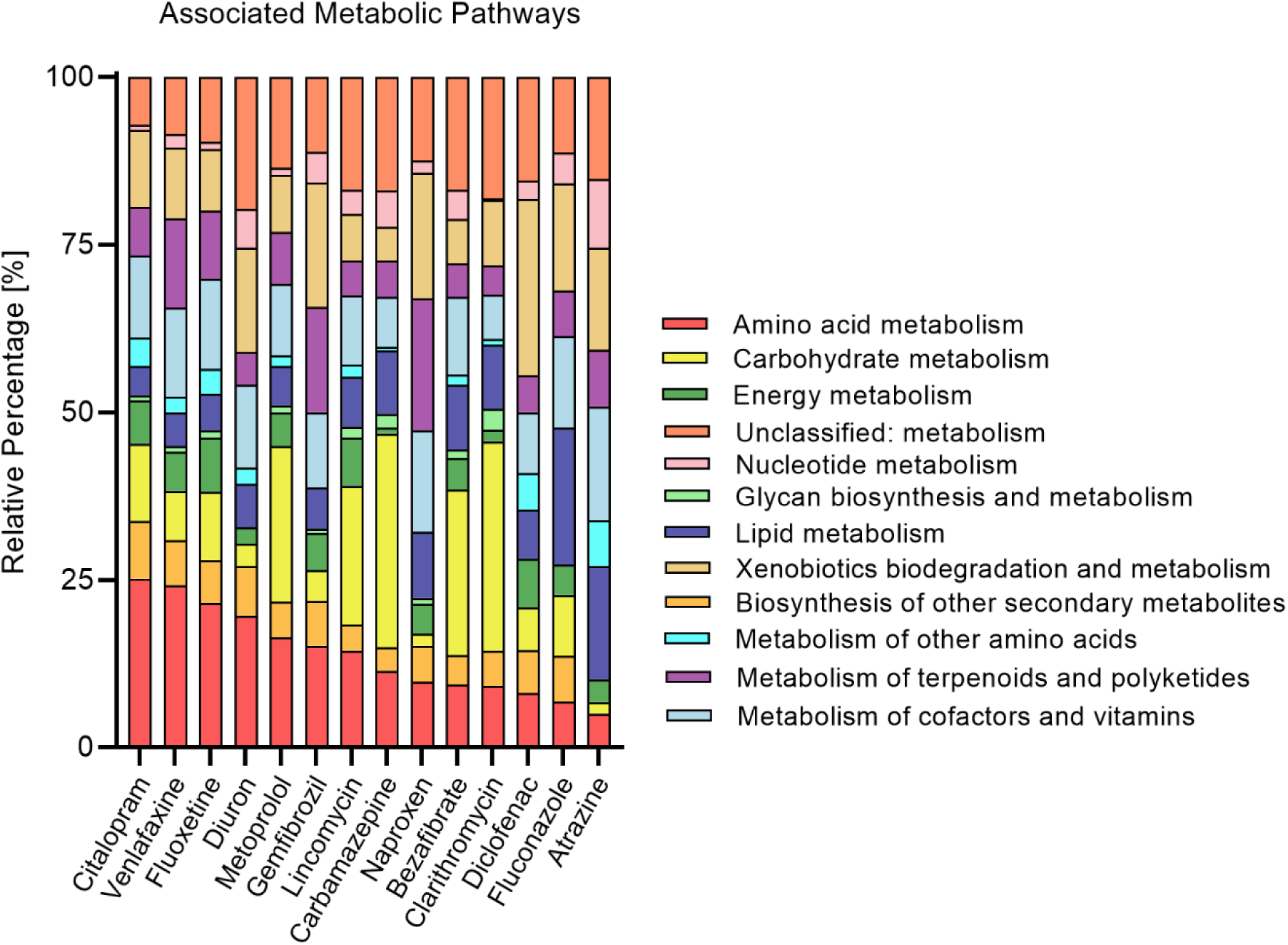
Metabolic origins of genes associated with OMP degradation. Bar chart displaying the relative percentage of genes from various microbial metabolic pathways potentially involved in the degradation of individual OMPs.

Amino acid metabolism emerged as particularly important for the degradation of amine-containing OMPs. Our correlation analysis showed that OMPs such as fluoxetine, venlafaxine, citalopram, and metoprolol strongly correlated with genes (enzymes) involved in amino acid metabolism (Figure 4). Furthermore, our results revealed a significant accumulation of free amino acids under conditions B and C (Figure S5), coinciding with enhanced degradation of amine-containing OMPs. Specifically, we detected 14 intracellular amino acids, including asparagine, glutamic acid, glycine, lysine, proline, serine, tryptophan, tyrosine, and valine, with concentrations significantly higher in condition C compared to condition A (ANOVA, P<0.05) (Figure S5). This accumulation of amino acids aligns with the significantly higher removal efficiency observed for fluoxetine (9 ± 3%), citalopram (9 ± 3%), venlafaxine (11 ± 2%), and metoprolol (35 ± 4%) under condition C (ANOVA, P<0.05, Figure 1). These findings suggest that the cyclic aeration in condition C may stimulate amino acid metabolism, potentially enhancing the co-metabolic degradation of amine-containing OMPs.

Our analysis also indicated a potential role for lipid metabolism in the degradation of OMPs containing carboxylic acid groups (-COOH) like bezafibrate and naproxen (Figure 4). Examining fatty acid profiles in our metabolomics data, we observed a significant increase in the abundance of long-chain fatty acids, including myristic acid, palmitoleic acid, linoleic acid, arachidic acid, and behenic acid under conditions C over condition A (Figure S5). This increase in fatty acid concentrations aligns with the enhanced degradation of bezafibrate (6 ± 2%) and naproxen (7 ± 3%) observed under the same conditions (Figure 1), suggesting that stimulating lipid metabolism, especially long-chain fatty acid accumulation under cyclic aeration conditions, may contribute to enhanced degradation.

Additionally, our correlation analysis suggested that xenobiotics biodegradation pathways might also be important for degrading structurally diverse OMPs such as gemfibrozil, diclofenac, and diuron (Figure 4, Table S6). While our metabolomics data does not directly measure xenobiotic metabolism intermediates, the overall increase in metabolic activity observed under conditions B and C, evidenced by the accumulation of amino acids, fatty acids, and other metabolites, suggests a general upregulation of diverse metabolic processes. This could include an increase in OMP co-metabolism, supporting our observation. These findings collectively suggest that cyclic aeration conditions (B and C) stimulate a broader range of metabolic processes, potentially enhancing the co-metabolic degradation of a diverse array of OMPs. Notably, these associations have not been previously reported, and future studies could confirm the involvement of these metabolic pathways in specific OMP degradation and assess the application of cyclic aeration in scaling up WWTP processes. We suggest future studies apply LC-MS-based metabolomics along with reactor operations to verify global changes in metabolome. Isotope-labeled studies could trace the fate of OMPs through these metabolic pathways, providing more direct evidence of their involvement in OMP degradation. Lastly, it is important to note that while these metabolomics results support our observations, the diversity within the sludge microbiome should be evaluated simultaneously to provide a more complete picture of the degradation processes.

### 3.7 Linking OMP degradation with key enzymes and microbial contributors

Despite extensive research, our understanding of how diverse microbes break down structurally different OMPs in WWTPs remains incomplete (Kennes-Veiga et al., 2022a). To address this gap, we propose a comprehensive framework that considers community members and elucidates the connections between OMP structural features, potential degradation genes, key metabolic functions, and microbial hosts involved in OMP degradation. Figure 5 illustrates the links between each OMP and potential EC sub-subclasses via biotransformation rules (btrules) and the microbial phyla associated with each EC sub-subclass, identified through multi-omics analysis. We categorize the networks based on OMP degradation efficiency: poor degradation (<40%), moderate degradation (40-70%), and high degradation (70-95%). This categorization highlights the different structural features, enzyme sub-subclasses, and microbial hosts that could contribute to the degradation (Table S7-9).

**Figure 5.**
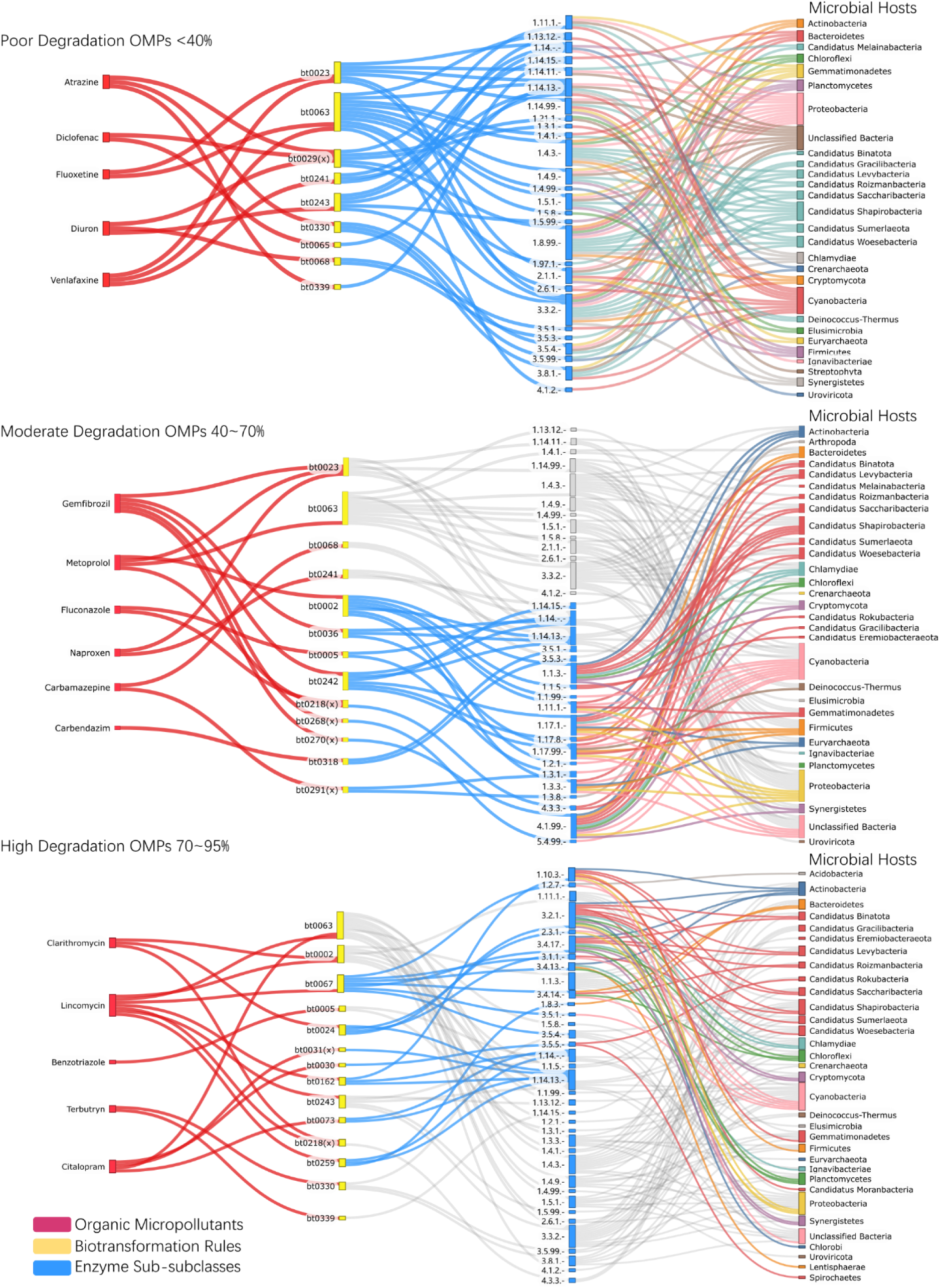
Network analysis of OMP co-metabolic degradation pathways. Connections involve OMP structures, biotransformation rules, enzyme sub-subclasses, and bacterial phyla. OMPs are categorized by degradation efficiency: poor (<40%), moderate (40-70%), and high (70-95%) removal. Highlighted connections in moderate and high removal categories represent unique pathways not observed in poorly degraded OMPs. Linkages between enzyme sub-subclasses and bacterial phyla are based on Mantel test results (P<0.05, Rho>0.5).

Our analysis reveals that OMPs with poor degradation rates (<40%) often exhibit less biodegradable structural features, posing challenges to their effective removal. However, we identified various peroxidases (EC 1.11.1.-) that could facilitate the removal of these persistent OMPs through the free radical mechanism (Aboelnga, 2022), which holds promise for enhancing the removal of halogen-containing OMPs. Our metaproteomics analysis revealed the presence of 8 distinct types of peroxidases in our system (Figure S3). Future studies should focus on validating the role of peroxidases in degrading persistent OMPs through *in vitro* pure enzyme reactions.

Our results indicate that dominant bacterial phyla such as *Actinobacteria*, *Bacteroidete*, *Chloroflexi*, *Proteobacteria*, and *Nitrospirae* play crucial roles in OMP removal at trace levels (Fang et al., 2018; Gao et al., 2019; Harb et al., 2016). Notably, the microbial abundance from tested conditions was highly correlated with the accumulated gene captured under each EC sub-subclass, as obtained by Mantel tests (Rho>0.5, P<0.05). This aligns with previous studies demonstrating the involvement of *Actinobacteria* and its member *Corynebacterium* in the degradation of sulfamethoxazole in WWTP (Kennes-Veiga et al., 2022b). Our data suggests that each microbial phylum and the functional genes they harbor, based on their assigned EC categories, can target various functional groups present in structurally diverse OMPs (Figure 5). Notably, the catalytic efficiency of these enzymes towards similar functional groups on various OMPs (Table S3) can vary and require further *in vitro* degradation studies to elucidate.

To better understand the complex co-metabolic degradation processes in WWTPs and inform strategies for targeted OMP removal, we developed a data analysis pipeline (Figure S2) that constrains omics datasets using both experimental OMP degradations and *in-silico* predictions based on the functional groups in the OMP structure. For amine-containing compounds undergoing potential dealkylations (bt0063) (Figure 5), we identified *Cyanobacteria* with strong (P<0.05, Rho>0.6) linkage with monooxygenase (EC:1.13.12.-), *Bacteroidetes* process strong correlations (Rho>0.6, P<0.05) with dehydrogenase (EC: 1.4.1.-), and aminotransferase (EC:2.6.1.-) was found to be most correlated with *Cyanobacteria* and *Proteobacteria* (P<0.05). These bacterial members could contribute significantly to our system’s overall degradation of amine-containing OMPs. We encourage future studies to adopt a similar multi-omics approach with different wastewater communities on-site to verify and generalize these linkages, ultimately leading to more effective strategies for OMP removal in WWTPs

## 4. Conclusion

This study presents an integrated multi-omics approach to elucidate the complex networks governing OMP co-metabolic degradation with wastewater microbiomes. We propose key enzyme candidates of oxidoreductases, peroxidases, hydrolases, and cytochrome P450s, metabolic pathways like amino acid, lipid, and xenobiotic metabolism, originated from microbial phyla of *Actinobacteria*, *Bacteroidetes*, and *Proteobacteria* are critical for the co-metabolic degradation of functional groups of halogen (-Cl, -F), amide (-CONH₂), amine (-NH₂) and carboxylic acid (-COOH) readily identified on various OMPs. We establish connections between OMP functional groups, degradation enzymes, metabolic pathways, and microbial phyla, providing a comprehensive framework for understanding OMP co-metabolic degradation pathways. It is important to note that the correlative hypothesis does not prove a direct catalytic mechanism for OMP degradation but a direction for future studies to prioritize. Future research should include heterologous expression of identified enzymes to examine the detailed reaction mechanism and isotope labeling studies could provide direct evidence of specific metabolic pathway involvement. We acknowledge the limitations of the controlled laboratory setting with a limited number of OMPs. Future studies should cope with different treatment parameters in WWTPs and additional OMPs covering more diverse structures. Despite these limitations, the multi-omics pipeline can be applied to other wastewater systems to generalize the findings and inform the development of predictive models for OMP fate. Our analysis suggests that cyclic aeration conditions could potentially enhance OMP removal in full-scale WWTPs. The identified enzyme candidates and metabolic pathways provide valuable targets for bioengineering approaches to improve OMP degradation efficiency. In conclusion, this study significantly advances our understanding of OMP co-metabolic degradation. It paves the way for developing more targeted and effective strategies for OMP removal in wastewater treatment systems, contributing to improved water quality and environmental protection.

## Abbreviations

OMPs: Organic micropollutants
WWTPs: Wastewater treatment plants
DO: Dissolved oxygen
SPE: Solid phase extraction
ECs: Enzyme commission numbers
KEGG: Kyoto Encyclopedia of Genes and Genomes
KO IDs: KEGG ortholog IDs
MLSS: Mixed liquor suspended solids
Eawag-PPD: Eawag Pathway Prediction System
enviPath: Environmental Contaminant Biotransformation Pathway Resource

## Acknowledgments

We thank Emma Jay for assisting with reactor operation and degradation analysis. We thank Watercare Services Ltd for providing the activated sludge culture. The authors acknowledge the use of New Zealand eScience Infrastructure (NeSI) high performance computing facilities, consulting support and/or training services as part of this research. The authors also acknowledge the Centre for eResearch at the University of Auckland for their help in facilitating this research.

## Supporting Information

All the supplementary tables and figures are provided in Supporting Information.

