## Supplemental Material for "Microbial Metabolic Enzymes, Pathways and Microbial Hosts for Co-Metabolic Degradation of Organic Micropollutants in Wastewater"

Total Pages: 25

Text: 1

Tables: 9

Figures: 5

**Supplementary Text**

**Text 1 Metagenomics analysis pipeline.**

DNA extraction from sludge samples was performed using the DNeasy PowerSoil Pro Kit (QIAGEN, Germany) according to the standard protocol. The downstream metagenomics process was outsourced to the Auckland Genomic Center (Auckland, NZ) with steps include DNA concentration, quality assessment of the DNA purity, and preparation for PCR and indexing. After post-library quality control (QC), normalization was performed, and a final pool check was conducted using a bioanalyzer. The samples were sequenced using the HiSeqX platform with 2×150 bp paired-end shotgun sequencing. We performed preprocessing steps to eliminate low-quality sequences using Trimmomatic (Bolger et al., 2014) with specific quality thresholds: LEADING:3, TRAILING:3, SLIDINGWINDOW:10:15, and MINLEN:50 with the following settings: TruSeq3-PE-2.fa:2:30:10:2:keepBothReads (Raes et al., 2021). For metagenomic taxonomic and functional profiling, we employed the SqueezeMeta v1.5.2 pipeline (Tamames and Puente-Sánchez, 2019). Co-assembly was performed using Megahit (Li et al., 2015), and short contigs (<200 bps) were filtered out using prinseq (Schmieder and Edwards, 2011). Within the SqueezeMeta pipeline, Barrnap (Seemann, 2014) tool was used for predicting RNAs, while Prodigal (Hyatt et al., 2010) was utilized for predicting ORFs. Taxonomic ranks were assigned against NCBI GenBank nr database (Clark et al., 2016) with identify thresholds of 85, 60, 55, 50, 46, 42, and 40% for species, genus, family, order, class, phylum, and superkingdom ranks, respectively (Luo et al., 2014). Kyoto Encyclopedia of Genes and Genomes (Kanehisa and Goto, 2000), was used for functional assignments with Diamond (Buchfink et al., 2014) with a maximum e-value threshold of 1e-03 and sequence identify threshold of 50%. Raw metagenomes are available at European Nucleotide Archive with data identifier PRJEB74089.

**Supplementary Tables**

**Table S1** Trace elements and synthetic wastewater provided to the reactor

| **Chemical** | **Concentration (g/L)** |
| --- | --- |
| EDTA | 2.50 |
| ZnSO_4_·7H_2_O | 1.10 |
| CoCl_2_·6H_2_O | 0.80 |
| MnCl_2_·4H_2_O | 2.55 |
| MgSO_4_·7H_2_O | 20.0 |
| CuSO_4_·5H_2_O | 0.86 |
| (NH_4_)_6_Mo_7_O_24_·4H_2_O | 0.07 |
| CaCl_2_·2H_2_O | 2.75 |
| FeSO_4_·7H_2_O | 2.57 |
| **Synthetic wastewater** | **Concentration** |
| Methanol (mL/L) | 26.94 |
| NH_4_Cl (g/L) | 24.46 (C); 30.57 (B & A) |
| KH_2_PO_4_ (g/L) | 2.76 |
| K_2_HPO_4_ (g/L) | 2.76 |
| NaHCO_3_ (g/L) | 76.80 |

**Table S2.** Internal Standards used in this study

| **Chemical** | Formula |
| --- | --- |
| Acesulfame Potassium-^13^C_4_ | ^13^C_4_H_4_KNO_4_S |
| Atenolol-d_7_ | C₁₄H₁₅D₇N₂O₃ |
| Bezafibrate-d_4_ | C₁₉H₁₆D₄ClNO₄ |
| Clarithromycin-N-methyl-^13^C, D_3_ | C₃₇¹³CH₆₆D₃NO₁₃ |
| Diclofenac-D_4_ | C_14_H_7_D_4_Cl_2_NO_2_ |
| Fluconazole-D_4_ | C_13_H_8_D_4_F_2_N_6_O |
| Fluoxetine-D_5_ | C_17_D_5_H_13_F_3_NO |
| Metoprolol-D_7_ | C_15_H_18_D_7_NO_3_ |
| Sucralose-D_6_  Sulfamethoxazole-^13^C_6_ | C_12_H_13_D_6_Cl_3_O_8_  C_10_H_11_N_3_O_3_S |

**Table S3** OMPs with predicted Biotransformation rules and EC subclasses associated with each functional group

| OMPs | Structure | Btrules (EAWAG)  Functional Group Attacked | Corresponding ECs (Envi-Path), Predicted |
| --- | --- | --- | --- |
| Acetaminophen | 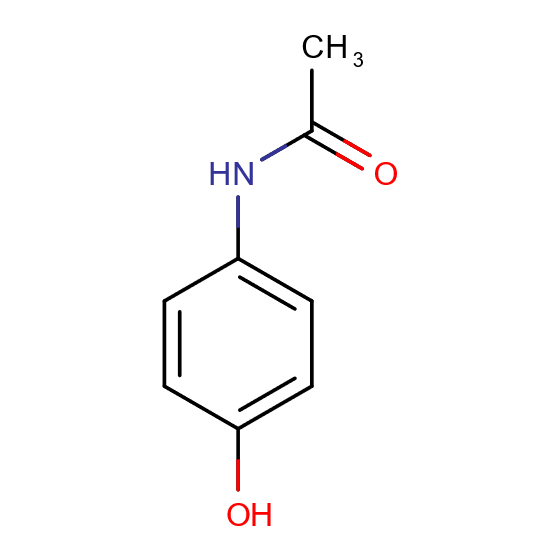 | bt0067 “secondary Amide to Carboxylate” | "1.2.7", "3.1.1", "3.4.13", "3.4.14", "3.4.17", "3.4.19", "3.5.1", "3.5.2", "3.5.4" |
| Atazanavir | 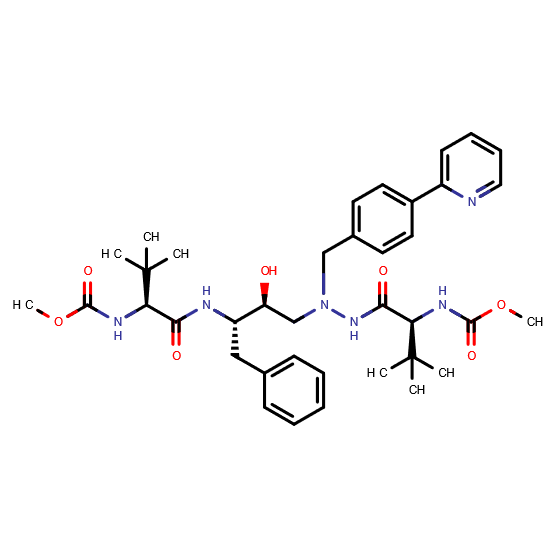 | bt0318 “Carbamyl to amine and carbonate” bt0002 “Secondary Alcohol Oxidation” bt0067 “secondary Amide to Carboxylate” bt0218 “2-Hydroxyethylamine Oxidation”(x) | "1.1.-", "1.1.1", "1.1.3", "1.1.5", "1.1.98", "1.1.99", "1.11.1", "1.14.12", "1.14.13", "1.2.1", "1.3.3", "1.2.7", "3.1.1",   "3.4.13", "3.4.14", "3.4.17", "3.4.19", "3.5.1", "3.5.2", "3.5.4", "3.5.3", "4.3.3" |
| Atrazine | 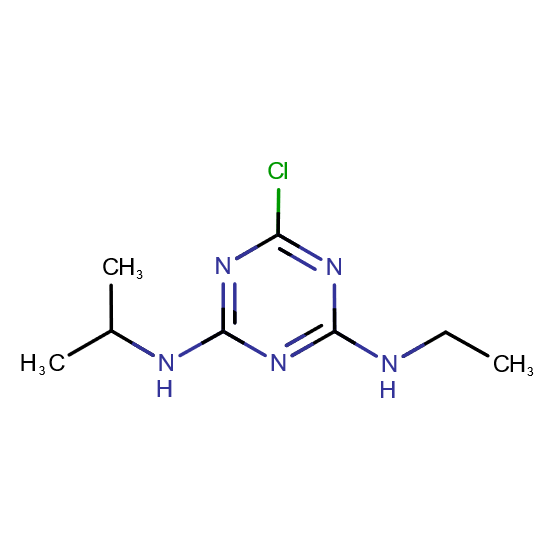 | bt0339 “N-aliphatic-s-triazine to Amino-s-triazine” bt0330 “1-halo-s-Triazine to 1-hydroxy-s-Triazine” bt0029 “Aliphatic dehalogenation”(x) | "1.14.15", "3.5.4", "3.5.99", "3.8.1", "1.11.1", "1.21.1", "1.3.1", "1.8.99", "1.97.1" |
| Benzotriazole | 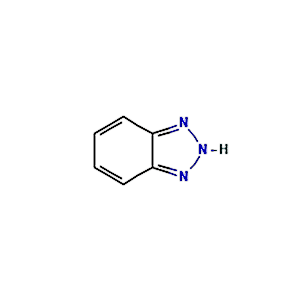 | bt0005 “Vic-unsubstituted Aromatic Dihydroxylation”* | "1.3.1", "1.14.-" |
| Bezafibrate | 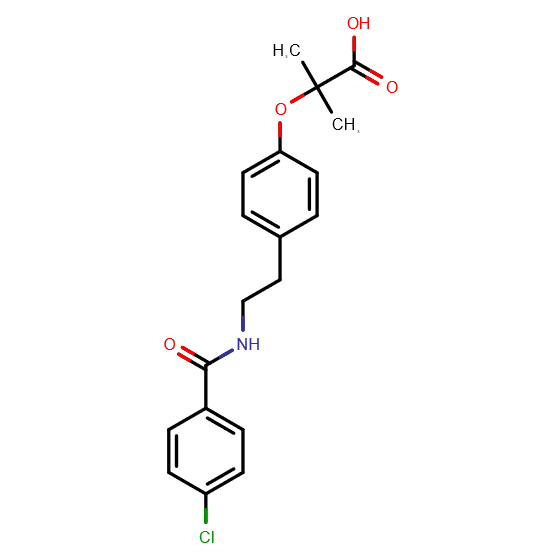 | bt0051 “2- or 3-substituted carboxylate” bt0243 “Amide Oxidation (Amide Dealkylation)” bt0067 “secondary Amide to Carboxylate” bt0029 “Aliphatic dehalogenation”(x) bt0268 “Carbon Chain Rearrangement”(x) | "2.1.3", "5.5.1", "1.1.1", "4.1.1", "7.2.4", "1.13.12", "1.14.13", "1.5.99", "3.3.2", "1.14.-", "1.2.7", "3.1.1",   "3.4.13", "3.4.14", "3.4.17", "3.4.19", "3.5.1", "3.5.2", "3.5.4", "1.11.1", "1.21.1", "1.3.1", "1.8.99", "1.97.1", "5.4.99" |
| Carbamazepine | 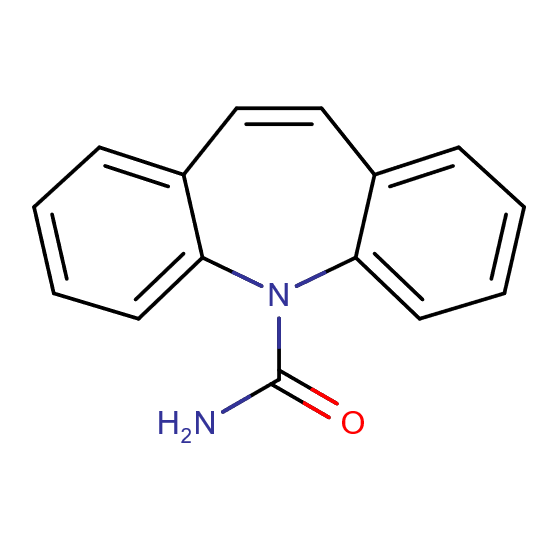 | bt0068 “Linear Urea Derivatives Hydroxylation “ bt0291 “Hydration of a double bond”(x) | "1.1.1", "1.3.1", "1.3.8", "3.5.1", "3.5.3" |
| Carbendazim | 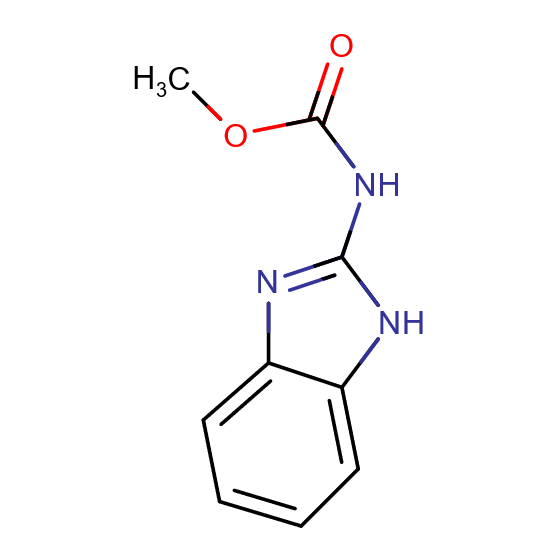 | bt0318 “Carbamyl to amine and carbonate”* | "3.5.1", "3.5.3" |
| Citalopram | 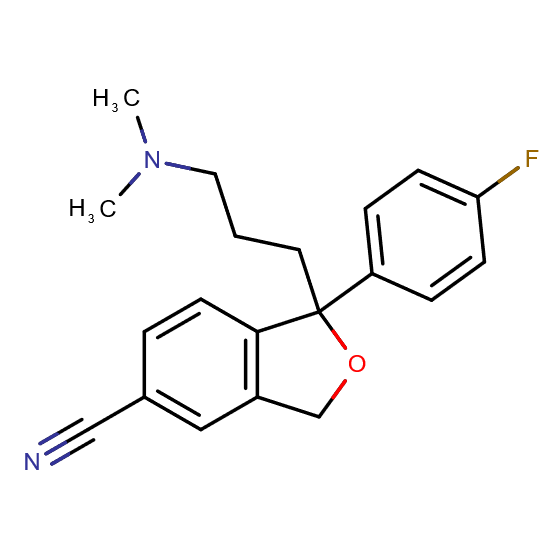 | bt0063 “Amine Oxidation (Amine Dealkylation)” bt0030 “Hydrolysis of Nitrile” bt0073 “UInsubstituted Cyclic ether to 2-Hydroxy cyclic ether” bt0031 “Benzonitrile Oxidation”(x) | "4.1.2", "1.13.12", "1.14.-", "1.14.13", "1.14.14", "1.4.1", "1.4.3", "1.4.9", "1.4.99", "1.5.1", "1.5.3", "1.5.8", "2.6.1", "3.3.2", "3.5.5" |
| Clarithromycin | 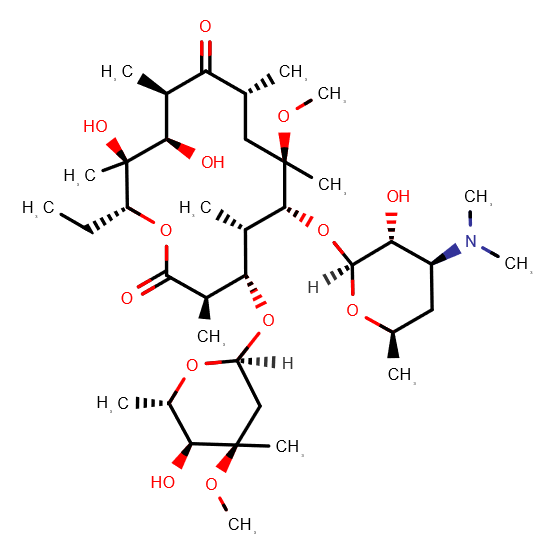 | bt0024 “Ester hydrolyses” bt0002 “Secondary Alcohol Oxidation” bt0218 “2-Hydroxyethylamine Oxidation”(x) | "1.1.-", "1.1.1", "1.1.3", "1.1.5", "1.1.98", "1.1.99", "1.11.1", "1.14.12", "1.14.13", "1.2.1", "1.3.3", "2.3.1", "3.1.1", "3.2.1", "4.3.3" |
| Climbazole | 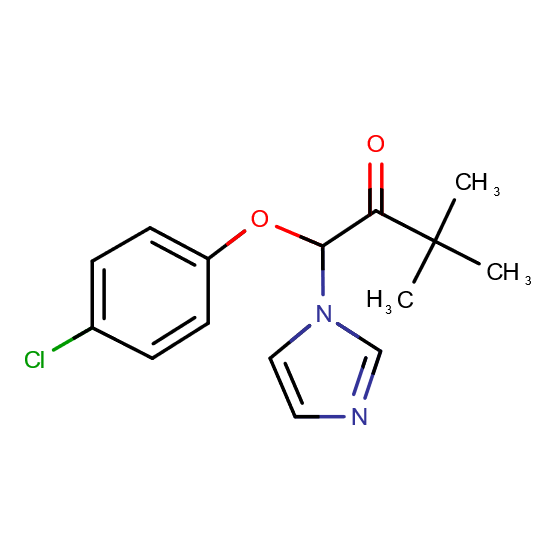 | bt0063 “Amine Oxidation (Amine Dealkylation)” bt0023 “Ether Oxidation” bt0029 “Aliphatic dehalogenation”(x) | "4.1.2", "1.13.12", "1.4.1", "1.4.3", "1.4.9", "1.4.99", "1.5.1", "1.5.3", "1.5.8", "2.6.1", "1.14.-",   "1.14.11", "1.14.13", "1.14.14", "1.14.99", "2.1.1", "3.3.2", "1.11.1", "1.21.1", "1.3.1", "1.8.99", "1.97.1" |
| Diclofenac | 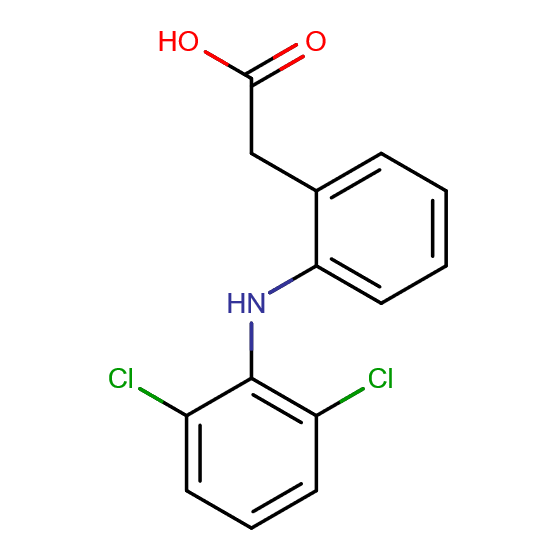 | bt0065 “Degradation of aromatic amines” bt0029 “Aliphatic dehalogenation”(x) | "1.14.12", "1.14.13", "1.11.1", "1.21.1", "1.3.1", "1.8.99", "1.97.1" |
| Diuron | 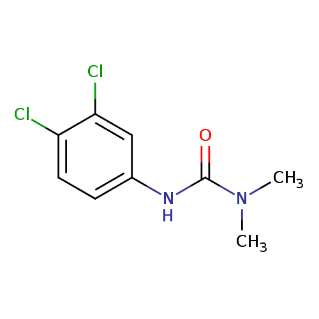 | bt0243 “Amide Oxidation (Amide Dealkylation)” bt0068 “Linear Urea Derivatives Hydroxylation “ bt0029 “Aliphatic dehalogenation”(x) | "1.13.12", "1.14.13", "1.5.99", "3.3.2", "1.14.-", "3.5.1", "3.5.3", "1.11.1", "1.21.1", "1.3.1", "1.8.99", "1.97.1" |
| Enrofloxacin | 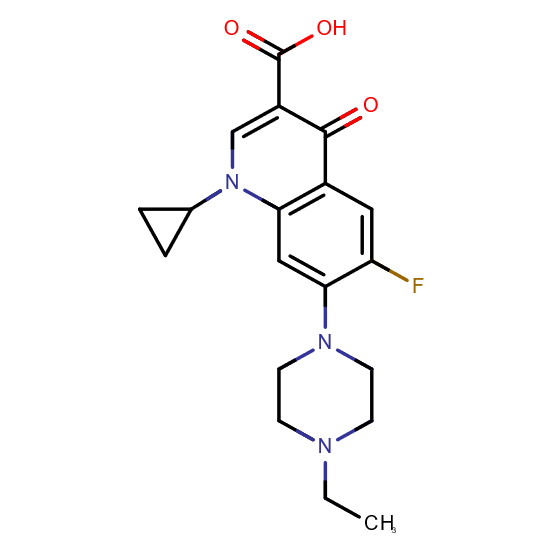 | bt0063 “Amine Oxidation (Amine Dealkylation)” | "4.1.2", "1.13.12", "1.14.13", "1.14.14", "1.4.1", "1.4.3", "1.4.9", "1.4.99", "1.5.1", "1.5.3", "1.5.8", "2.6.1", "3.3.2" |
| Estriol | 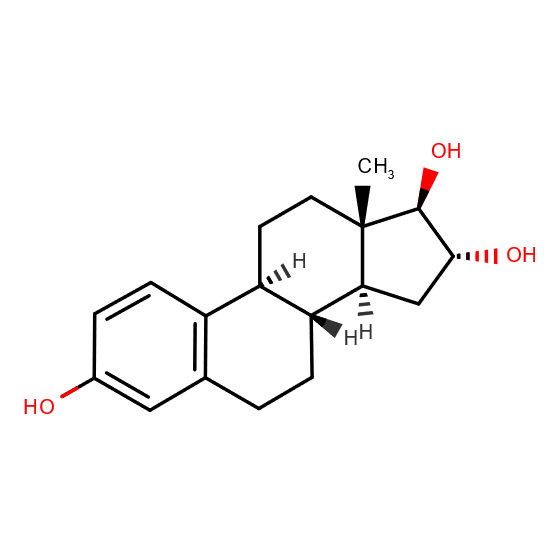 | bt0014 “1-hydroxy-2-unsubstituted aromatic” bt0002 “Secondary Alcohol Oxidation” | "1.14.-", "1.11.2", "1.13.11", "1.14.14", "1.14.18", "1.1.-", "1.1.1", "1.1.3", "1.1.5", "1.1.98", "1.1.99", "1.11.1", "1.14.12", "1.14.13", "1.2.1", "1.3.3" |
| Fluconazole | 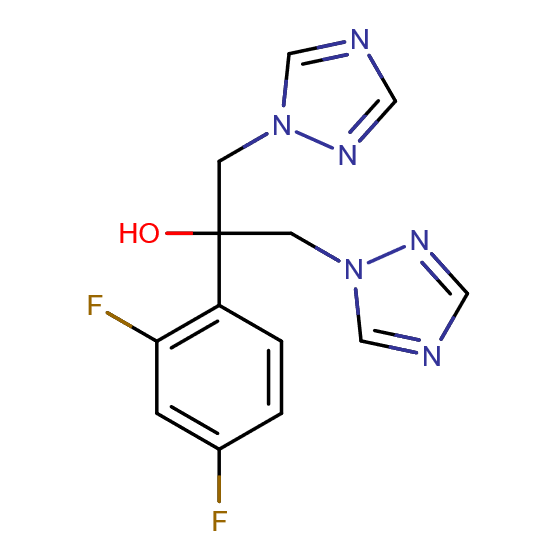 | bt0005 “Vic-unsubstituted Aromatic Dihydroxylation”* bt0218 “2-Hydroxyethylamine Oxidation”(x) | "1.3.1", "1.14.-", "4.3.3" |
| Fluoxetine | 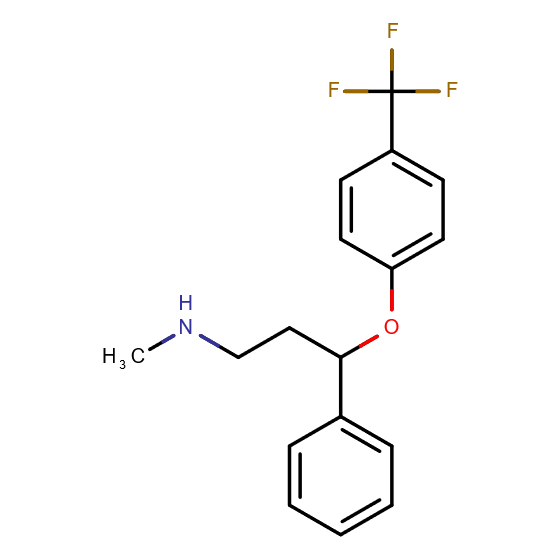 | bt0023 “Ether Oxidation” bt0063 “Amine Oxidation (Amine Dealkylation)” | "1.14.-", "1.14.11", "1.14.99", "2.1.1", "4.1.2", "1.13.12", "1.14.13", "1.14.14", "1.4.1", "1.4.3", "1.4.9", "1.4.99", "1.5.1", "1.5.3", "1.5.8", "2.6.1", "3.3.2" |
| Gemfibrozil | 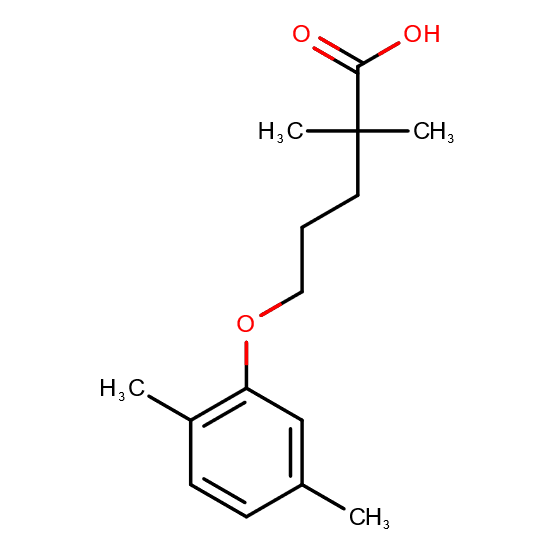 | bt0023 “Ether Oxidation” bt0036 “Aromatic Methyl Monohydroxylation” bt0242 “Secondary Aliphatic Monohydroxylation” bt0268 “Carbon Chain Rearrangement”(x) bt0270 “Methylsuccinate addition to aromatic methyl”(x) | "1.14.-", "1.14.11", "1.14.13", "1.14.14", "1.14.99", "2.1.1", "3.3.2", "1.14.12", "1.14.15", "1.17.1", "1.17.8", "1.17.99", "4.1.99", "5.4.99" |
| Lincomycin | 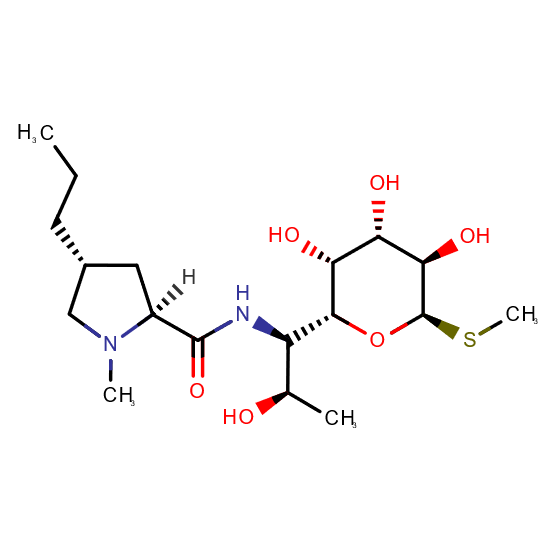 | bt0063 “Amine Oxidation (Amine Dealkylation)” bt0002 “Secondary Alcohol Oxidation” bt0067 “secondary Amide to Carboxylate” bt0162 “Sulfide to sulfoxide” bt0243 “Amide Oxidation (Amide Dealkylation)” bt0259 “Sulfide cleavages” bt0218 “2-Hydroxyethylamine Oxidation”(x) | "4.1.2", "1.13.12", "1.14.14", "1.4.1", "1.4.3", "1.4.9", "1.4.99", "1.5.1", "1.5.3", "1.5.8", "2.6.1", "3.3.2", "1.1.-", "1.1.3",   "1.1.5", "1.1.98", "1.1.99", "1.11.1", "1.14.12", "1.2.1", "1.3.3", "1.2.7", "3.1.1", "3.4.13", "3.4.14", "3.4.17", "3.4.19", "3.5.1",   "3.5.2", "3.5.4", "1.14.-", "1.10.3", "1.13.12", "1.14.13", "1.5.99", "1.1.1", "1.8.3", "4.3.3" |
| Metoprolol | 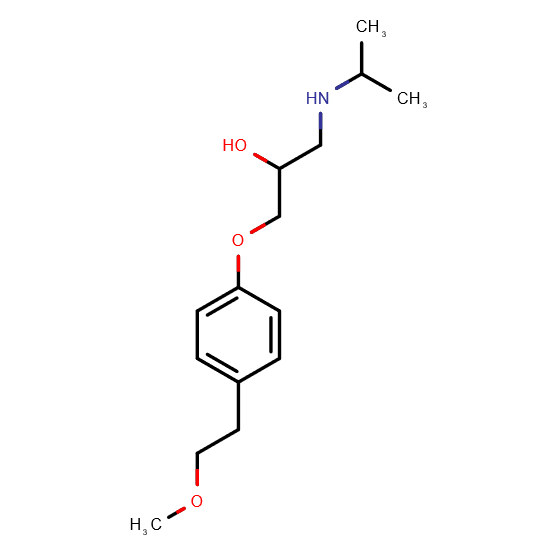 | bt0063 “Amine Oxidation (Amine Dealkylation)” bt0002 “Secondary Alcohol Oxidation” bt0023 “Ether Oxidation” bt0218 “2-Hydroxyethylamine Oxidation”(x) | "4.1.2", "1.13.12", "1.14.13", "1.14.14", "1.4.1", "1.4.3", "1.4.9", "1.4.99", "1.5.1", "1.5.3", "1.5.8", "2.6.1", "3.3.2", "1.1.-",   "1.1.1", "1.1.3", "1.1.5", "1.1.98", "1.1.99", "1.11.1", "1.14.12", "1.2.1", "1.3.3", "1.14.-", "1.14.11", "1.14.99", "2.1.1", "4.3.3" |
| Naproxen | 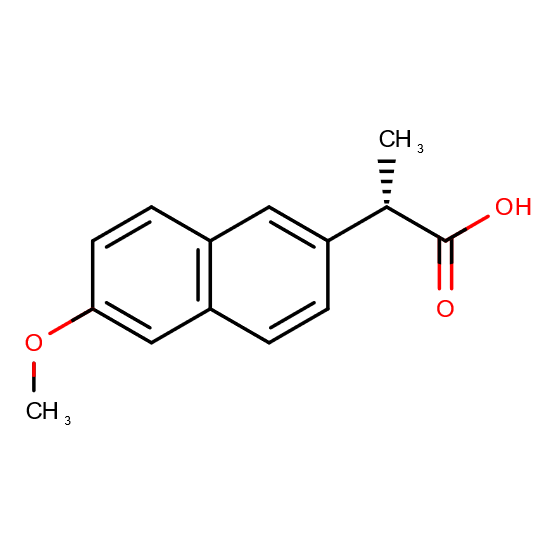 | bt0241 “Monohydroxylation” bt0023 “Ether Oxidation” | "1.14.-", "1.14.11", "1.14.13", "1.14.14", "1.14.15", "1.14.99", "2.1.1", "3.3.2" |
| Sulfamethoxazole | 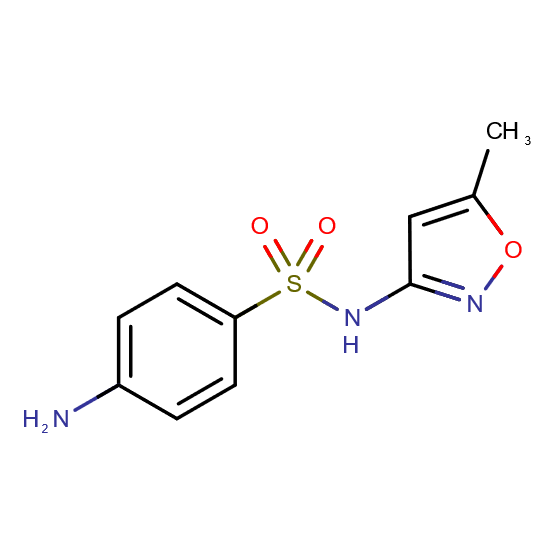 | bt0144 “Sulphonamides Hydrolysis” | "3.10.1" |
| Terbutryn | 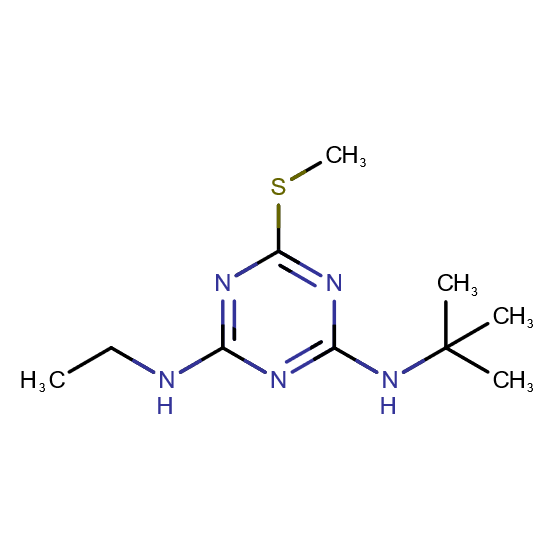 | bt0330 “1-halo-s-Triazine to 1-hydroxy-s-Triazine” bt0339 “*N*-aliphatic-s-triazine to Amino-s-triazine” | "1.14.15", "3.5.4", "3.5.99", "3.8.1" |
| Tylosin | 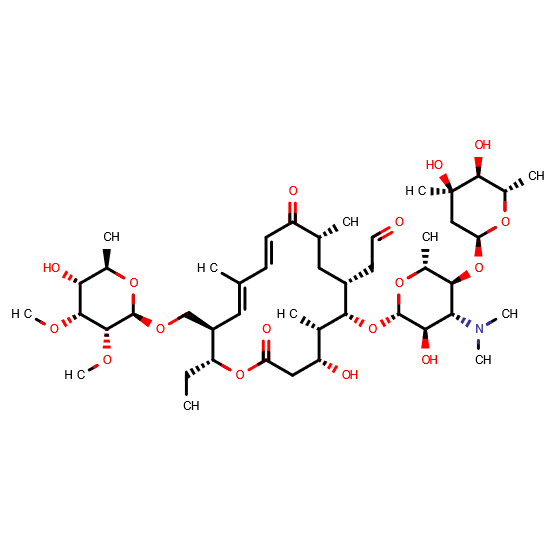 | bt0003 “Aldehyde -> Carboxylate” | "1.2.-", "4.1.2", "1.1.1", "1.1.3", "1.14.13", "1.2.1", "1.2.3" |
| Venlafaxine | 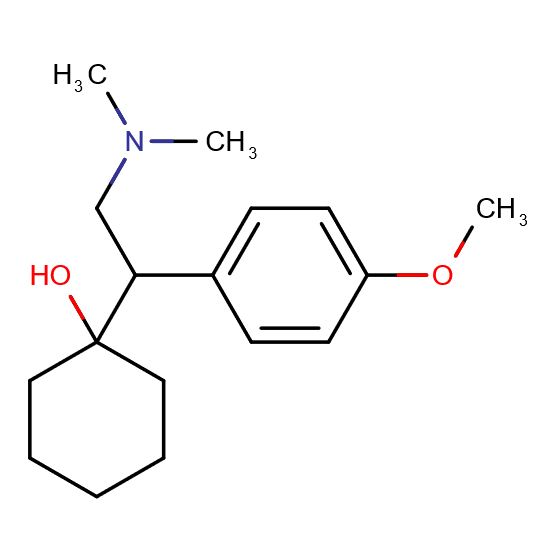 | bt0063 “Amine Oxidation (Amine Dealkylation)” bt0023 “Ether Oxidation” bt0241 “Monohydroxylation” | "4.1.2", "1.13.12", "1.14.13", "1.14.14", "1.4.1", "1.4.3", "1.4.9", "1.4.99", "1.5.1", "1.5.3", "1.5.8", "2.6.1", "1.14.11", "1.14.99", "2.1.1", "3.3.2", "1.14.-", "1.14.15" |

**Table S4** Metagenomics data detected EC subclasses and the number of different KEGG Orthology IDs under each EC sub-subclasses

| EC Subclasses | Number of KO IDs detected | EC Subclasses | Number of KO IDs detected |
| --- | --- | --- | --- |
| 1.1.-.-; 1.1.1.-; 1.1.3.-; 1.1.5.-; 1.1.98.-; 1.1.99.-; 1.10.3.-; 1.11.1-; 1.11.2.-; 1.13.11.-; 1.13.12.-; 1.14.-.-; 1.14.11.-; 1.14.12.-; 1.14.13.-; 1.14.14.-; 1.14.15.-; 1.14.18.-; 1.14.99.-; 1.17.1.-; 1.17.8.-; 1.17.99.-; 1.2.-.-; 1.2.1.-; 1.2.3.-; 1.2.7.-; 1.21.1.-; 1.3.1.-; 1.3.3.-; 1.3.8.-; 1.4.1.-; 1.4.3.-; 1.4.9.-; 1.4.99.-; 1.5.1.-; 1.5.3.-; 1.5.8.-; 1.5.99.-; 1.8.3.-; 1.8.99.-; 1.97.1.-; 2.1.1.-; 2.1.3.-; 2.3.1.-; 2.6.1.-; 3.1.1.-; 3.10.1.-; 3.2.1.- | 11; 252; 17; 13; 3; 15; 6; 25; 3; 47; 3; 11; 28; 19; 81; 33; 23; 6; 15; 20; 1; 4; 1; 64; 4; 24; 1; 68; 5; 14; 17; 12; 3; 3; 31; 13; 3; 14; 4; 4; 8; 199; 10; 171; 80; 76; 1; 103 | 3.3.2.-; 3.4.13.-; 3.4.14.-; 3.4.17.-; 3.4.19.-; 3.5.1.-; 3.5.2.-; 3.5.3.-; 3.5.4.-; 3.5.5.-; 3.5.99.-; 3.8.1.-; 4.1.1.-; 4.1.2.-; 4.1.99.-; 4.3.3.-; 5.4.99.-; 5.5.1.-; 7.2.4.- | 9 \|; 10 \|; 10 \|; 18 \|; 11 \|; 103 \|; 30 \|; 16 \|; 30 \|; 6 \|; 7 \|; 3 \|; 81 \|; 39 \|; 13 \|; 4 \|; 36 \|; 9 \|; 5 \| |

*

**EC: 1. -. -.- Oxidoreductases.**

**EC: 1. 1. -.-	Acting on the CH-OH group of donors.**

**EC: 1. 2. -.-	Acting on the aldehyde or oxo group of donors.**

**EC: 1. 3. -.-	Acting on the CH-CH group of donors.**

**EC: 1. 4. -.-	Acting on the CH-NH2 group of donors.**

**EC: 1. 5. -.-	Acting on the CH-NH group of donors.**

**EC: 1. 8. -.-	Acting on a sulfur group of donors.**

**EC: 1.10. -.-	Acting on diphenols and related substances as donors.**

**EC: 1.11. -.-	Acting on a peroxide as acceptor.**

**EC: 1.13. -.- Acting on single donors with incorporation of molecular oxygen (oxygenases). The oxygen incorporated need not be derived from O2.**

**EC: 1.14. -.- Acting on paired donors, with incorporation or reduction of molecular oxygen. The oxygen incorporated need not be derived from O2.**

**EC: 1.17. -.-	Acting on CH or CH2 groups.**

**EC: 1.21. -.-	Catalyzing the reaction X-H + Y-H = 'X-Y'.**

**EC: 1.97. -.-	Other oxidoreductases.**

**EC: 2. -. -.- Transferases.**

**EC: 2. 1. -.-	Transferring one-carbon groups.**

**EC: 3. -. -.- Hydrolases.**

**EC: 3. 3. -.-	Acting on ether bonds.**

**EC: 3. 4. -.-	Acting on peptide bonds (peptidases).**

**EC: 3. 5. -.-	Acting on carbon-nitrogen bonds, other than peptide bonds.**

**EC: 3. 8. -.-	Acting on halide bonds.**

**EC: 4. -. -.- Lyases.**

**EC: 4. 1. -.-	Carbon-carbon lyases.**

**EC: 4. 3. -.-	Carbon-nitrogen lyases.**

**EC: 5. -. -.- Isomerases.**

**EC: 5. 4. -.-	Intramolecular transferases.**

**EC: 5. 5. -.-	Intramolecular lyases.**

**EC: 7. -. -.- Translocases.**

**EC: 7. 2. -.-	Catalysing the translocation of inorganic cations.**

**Table S5** Number of KO IDs assigned under each biotransformation rule

| Btrules | Number of KO IDs |
| --- | --- |
| bt0002 “Secondary Alcohol Oxidation” | 505 |
| bt0003 “Aldehyde -> Carboxylate” | 458 |
| bt0005 “Vic-unsubstituted Aromatic Dihydroxylation” | 79 |
| bt0014 “1-hydroxy-2-unsubstituted aromatic” | 181 |
| bt0023 “Ether Oxidation” | 376 |
| bt0024 “Ester hydrolyses” | 431 |
| bt0029 “Aliphatic dehalogenation”(x) | 106 |
| bt0030 “Hydrolysis of Nitrile” | 6 |
| bt0031 “Benzonitrile Oxidation”(x) | 81 |
| bt0036 “Aromatic Methyl Monohydroxylation” | 96 |
| bt0051 “2- or 3-substituted carboxylate” | 433 |
| bt0063 “Amine Oxidation (Amine Dealkylation)” | 327 |
| bt0065 “Degradation of aromatic amines” | 100 |
| bt0067 “secondary Amide to Carboxylate” | 312 |
| bt0068 “Linear Urea Derivatives Hydroxylation “ | 119 |
| bt0073 “UInsubstituted Cyclic ether to 2-Hydroxy cyclic ether” | 92 |
| bt0144 “Sulphonamides Hydrolysis” | 1 |
| bt0162 “Sulfide to sulfoxide” | 98 |
| bt0218 “2-Hydroxyethylamine Oxidation”(x) | 4 |
| bt0241 “Monohydroxylation” | 148 |
| bt0242 “Secondary Aliphatic Monohydroxylation” | 192 |
| bt0243 “Amide Oxidation (Amide Dealkylation)” | 118 |
| bt0259 “Sulfide cleavages” | 348 |
| bt0268 “Carbon Chain Rearrangement”(x) | 36 |
| bt0270 “Methylsuccinate addition to aromatic methyl”(x) | 13 |
| bt0291 “Hydration of a double bond”(x) | 334 |
| bt0318 “Carbamyl to amine and carbonate” | 119 |
| bt0330 “1-halo-s-Triazine to 1-hydroxy-s-Triazine” | 40 |
| bt0339 “N-aliphatic-s-triazine to Amino-s-triazine” | 23 |

**Supplementary Table S6.** Positively correlated genes under metabolic functions (KEGG Category level 2) associated with each OMP.

|  | Atrazine | Bezafibrate | Carbamazepine | Citalopram | Clarithromycin | Diclofenac | Diuron | Fluconazole | Fluoxetine | Gemfibrozil | Lincomycin | Metoprolol | Naproxen | Venlafaxine |
| --- | --- | --- | --- | --- | --- | --- | --- | --- | --- | --- | --- | --- | --- | --- |
| Amino acid metabolism | 3 | 30 | 23 | 35 | 42 | 9 | 24 | 3 | 40 | 27 | 56 | 62 | 11 | 62 |
| Biosynthesis of other secondary metabolites | 0 | 14 | 7 | 12 | 24 | 7 | 9 | 3 | 12 | 12 | 15 | 20 | 6 | 17 |
| Carbohydrate metabolism | 1 | 79 | 64 | 16 | 143 | 7 | 4 | 4 | 19 | 8 | 80 | 87 | 2 | 19 |
| Energy metabolism | 2 | 15 | 2 | 9 | 8 | 8 | 3 | 2 | 15 | 10 | 28 | 19 | 5 | 15 |
| Glycan biosynthesis and metabolism | 0 | 4 | 4 | 1 | 14 | 0 | 0 | 0 | 2 | 1 | 6 | 4 | 1 | 2 |
| Lipid metabolism | 10 | 31 | 19 | 6 | 44 | 8 | 8 | 9 | 10 | 11 | 29 | 22 | 11 | 13 |
| Metabolism of cofactors and vitamins | 10 | 37 | 15 | 17 | 30 | 10 | 15 | 6 | 25 | 20 | 40 | 40 | 17 | 34 |
| Metabolism of other amino acids | 4 | 5 | 1 | 6 | 4 | 6 | 3 | 0 | 7 | 0 | 7 | 6 | 0 | 6 |
| Metabolism of terpenoids and polyketides | 5 | 16 | 11 | 10 | 20 | 6 | 6 | 3 | 19 | 28 | 20 | 29 | 22 | 34 |
| Nucleotide metabolism | 6 | 14 | 11 | 1 | 1 | 3 | 7 | 2 | 2 | 8 | 14 | 4 | 2 | 5 |
| Unclassified: metabolism | 9 | 54 | 34 | 10 | 83 | 17 | 24 | 5 | 18 | 20 | 65 | 51 | 14 | 22 |
| Xenobiotics biodegradation and metabolism | 9 | 21 | 10 | 16 | 45 | 29 | 19 | 7 | 17 | 33 | 27 | 32 | 21 | 27 |

**Supplementary Table S7. Biotransformation rules associated with OMPs under different degradation groups.**

| Biotransformation Rules | | |
| --- | --- | --- |
| Low degradation <40% | Moderate degradation  40%-70% | High degradation 70-95% |
| bt0023 | bt0002 | bt0002 |
| bt0029(x) | bt0005 | bt0005 |
| bt0063 | bt0023 | bt0024 |
| bt0065 | bt0036 | bt0030 |
| bt0068 | bt0063 | bt0031(x) |
| bt0241 | bt0068 | bt0063 |
| bt0243 | bt0218(x) | bt0067 |
| bt0330 | bt0241 | bt0073 |
| bt0339 | bt0242 | bt0162 |
|  | bt0268(x) | bt0218(x) |
|  | bt0270(x) | bt0243 |
|  | bt0291(x) | bt0259 |
|  | bt0318 | bt0330 |
|  |  | bt0339 |

**Supplementary Table S8. Enzymes at sub-subclasses associated with OMPs under different degradation groups.**

| Enzymes at sub-subclasses | | |
| --- | --- | --- |
| Low degradation <40% | Moderate degradation  40%-70% | High degradation 70-95% |
| 1.11.1.- | 1.1.-.- | 1.1.-.- |
| 1.13.12.- | 1.1.1.- | 1.1.1.- |
| 1.14.-.- | 1.1.3.- | 1.1.3.- |
| 1.14.11.- | 1.1.5.- | 1.1.5.- |
| 1.14.12.- | 1.1.98.- | 1.1.98.- |
| 1.14.13.- | 1.1.99.- | 1.1.99.- |
| 1.14.14.- | 1.11.1.- | 1.10.3.- |
| 1.14.15.- | 1.13.12.- | 1.11.1.- |
| 1.14.99.- | 1.14.-.- | 1.13.12.- |
| 1.21.1.- | 1.14.11.- | 1.14.-.- |
| 1.3.1.- | 1.14.12.- | 1.14.12.- |
| 1.4.1.- | 1.14.13.- | 1.14.13.- |
| 1.4.3.- | 1.14.14.- | 1.14.14.- |
| 1.4.9.- | 1.14.15.- | 1.14.15.- |
| 1.4.99.- | 1.14.99.- | 1.2.1.- |
| 1.5.1.- | 1.17.1.- | 1.2.7.- |
| 1.5.3.- | 1.17.8.- | 1.3.1.- |
| 1.5.8.- | 1.17.99.- | 1.3.3.- |
| 1.5.99.- | 1.2.1.- | 1.4.1.- |
| 1.8.99.- | 1.3.1.- | 1.4.3.- |
| 1.97.1.- | 1.3.3.- | 1.4.9.- |
| 2.1.1.- | 1.3.8.- | 1.4.99.- |
| 2.6.1.- | 1.4.1.- | 1.5.1.- |
| 3.3.2.- | 1.4.3.- | 1.5.3.- |
| 3.5.1.- | 1.4.9.- | 1.5.8.- |
| 3.5.3.- | 1.4.99.- | 1.5.99.- |
| 3.5.4.- | 1.5.1.- | 1.8.3.- |
| 3.5.99.- | 1.5.3.- | 2.3.1.- |
| 3.8.1.- | 1.5.8.- | 2.6.1.- |
| 4.1.2.- | 2.1.1.- | 3.1.1.- |
|  | 2.6.1.- | 3.2.1.- |
|  | 3.3.2.- | 3.3.2.- |
|  | 3.5.1.- | 3.4.13.- |
|  | 3.5.3.- | 3.4.14.- |
|  | 4.1.2.- | 3.4.17.- |
|  | 4.1.99.- | 3.4.19.- |
|  | 4.3.3.- | 3.5.1.-/ 3.5.2.-/ 3.5.4.- |
|  | 5.4.99.- | 3.5.5.-/ 3.5.99.-/ 3.8.1.- |
|  |  | 4.1.2.-/ 4.3.3.- |

**Supplementary Table S9. Bacteria Phylum associated with OMPs under different degradation groups.**

| Bacteria Phylum | | |
| --- | --- | --- |
| Low degradation <40% | Moderate degradation  40%-70% | High degradation 70-95% |
| Actinobacteria | Bacteroidetes | Acidobacteria |
| Bacteroidetes | Candidatus Binatota | Actinobacteria |
| Candidatus Binatota | Candidatus Eremiobacteraeota | Bacteroidetes |
| Candidatus Gracilibacteria | Candidatus Gracilibacteria | Candidatus Binatota |
| Candidatus Levybacteria | Candidatus Levybacteria | Candidatus Eremiobacteraeota |
| Candidatus Melainabacteria | Candidatus Melainabacteria | Candidatus Gracilibacteria |
| Candidatus Roizmanbacteria | Candidatus Roizmanbacteria | Candidatus Levybacteria |
| Candidatus Saccharibacteria | Candidatus Rokubacteria | Candidatus Moranbacteria |
| Candidatus Shapirobacteria | Candidatus Saccharibacteria | Candidatus Roizmanbacteria |
| Candidatus Sumerlaeota | Candidatus Shapirobacteria | Candidatus Rokubacteria |
| Candidatus Woesebacteria | Candidatus Sumerlaeota | Candidatus Saccharibacteria |
| Chlamydiae | Candidatus Woesebacteria | Candidatus Shapirobacteria |
| Chloroflexi | Chlamydiae | Candidatus Sumerlaeota |
| Crenarchaeota | Chloroflexi | Candidatus Woesebacteria |
| Cryptomycota | Crenarchaeota | Chlamydiae |
| Cyanobacteria | Cryptomycota | Chlorobi |
| Deinococcus-Thermus | Cyanobacteria | Chloroflexi |
| Elusimicrobia | Deinococcus-Thermus | Crenarchaeota |
| Euryarchaeota | Elusimicrobia | Cryptomycota |
| Firmicutes | Euryarchaeota | Cyanobacteria |
| Gemmatimonadetes | Firmicutes | Deinococcus-Thermus |
| Ignavibacteriae | Gemmatimonadetes | Elusimicrobia |
| Planctomycetes | Ignavibacteriae | Euryarchaeota |
| Proteobacteria | Planctomycetes | Firmicutes |
| Streptophyta | Proteobacteria | Gemmatimonadetes |
| Synergistetes | Streptophyta | Ignavibacteriae |
| Unclassified Bacteria | Synergistetes | Lentisphaerae |
| Uroviricota | Unclassified Bacteria | Planctomycetes |
|  | Uroviricota | Proteobacteria |
|  |  | Spirochaetes |
|  |  | Streptophyta |
|  |  | Synergistetes |
|  |  | Unclassified Bacteria |
|  |  | Uroviricota |

**Supplementary Figures**

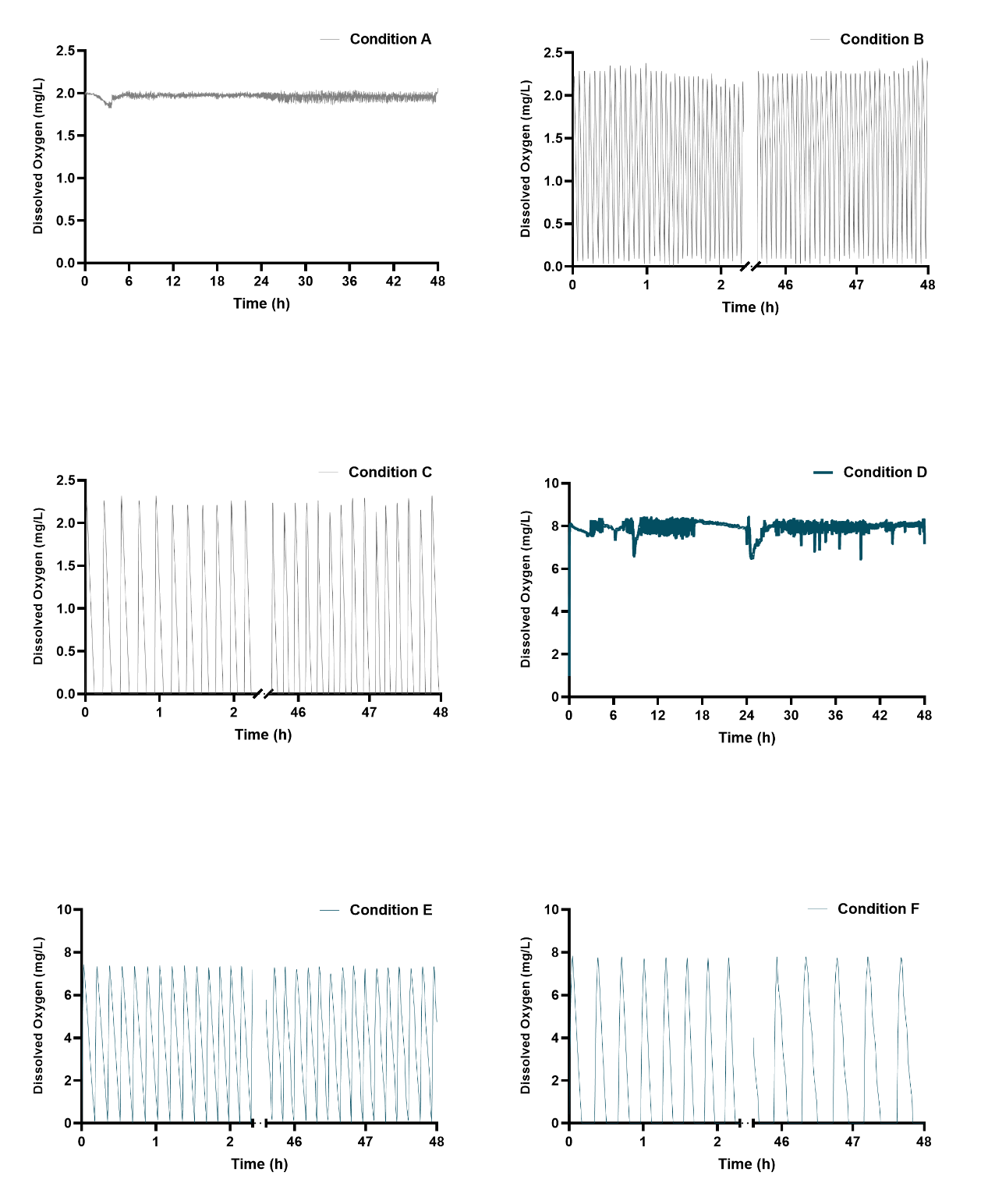

**Figure S1** Dissolved Oxygen Concentration of Conditions A, B, and C（Exp 1.）and D, E, and F (Exp 2.)

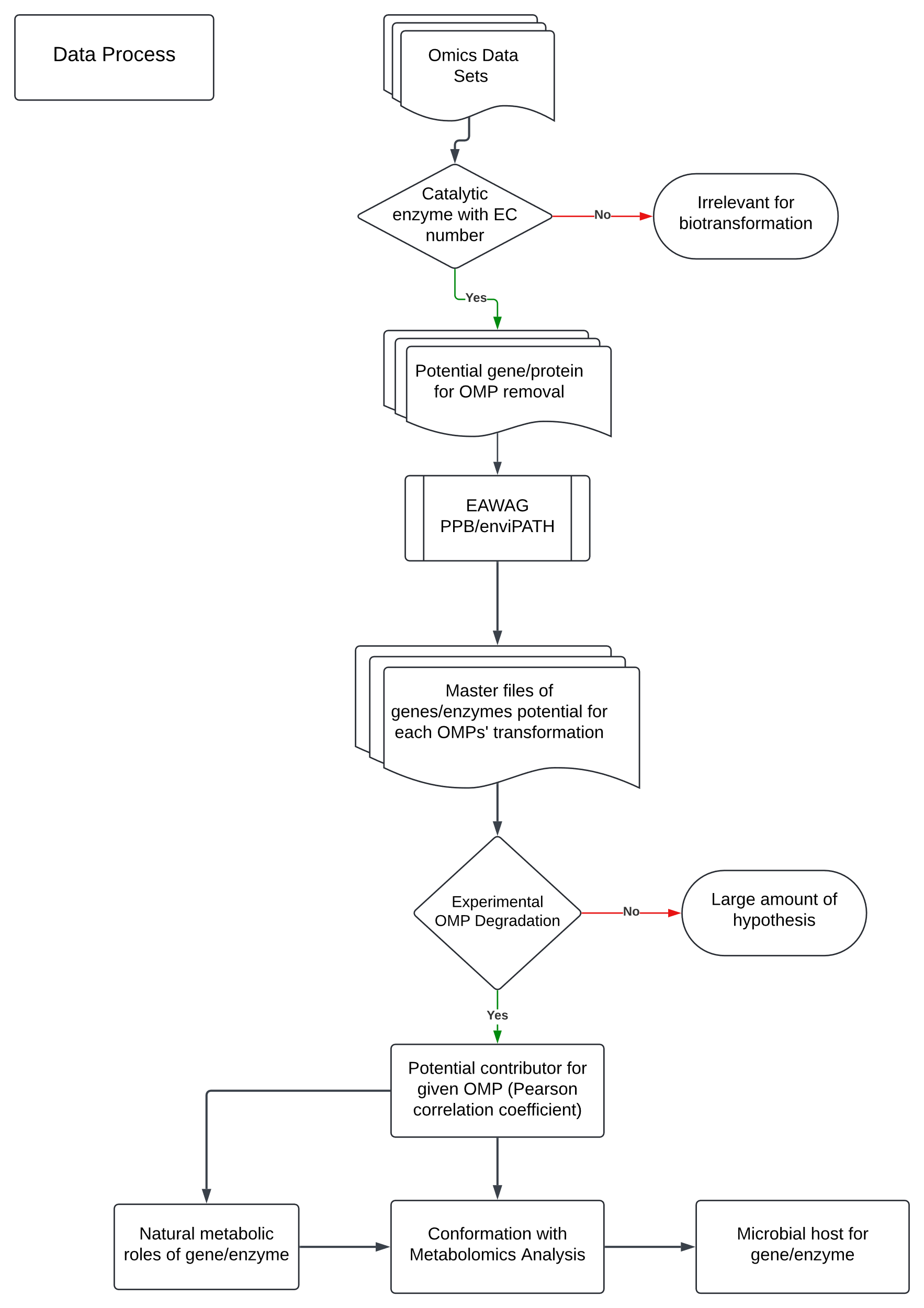

**Figure S2.** Roadmap for data processing

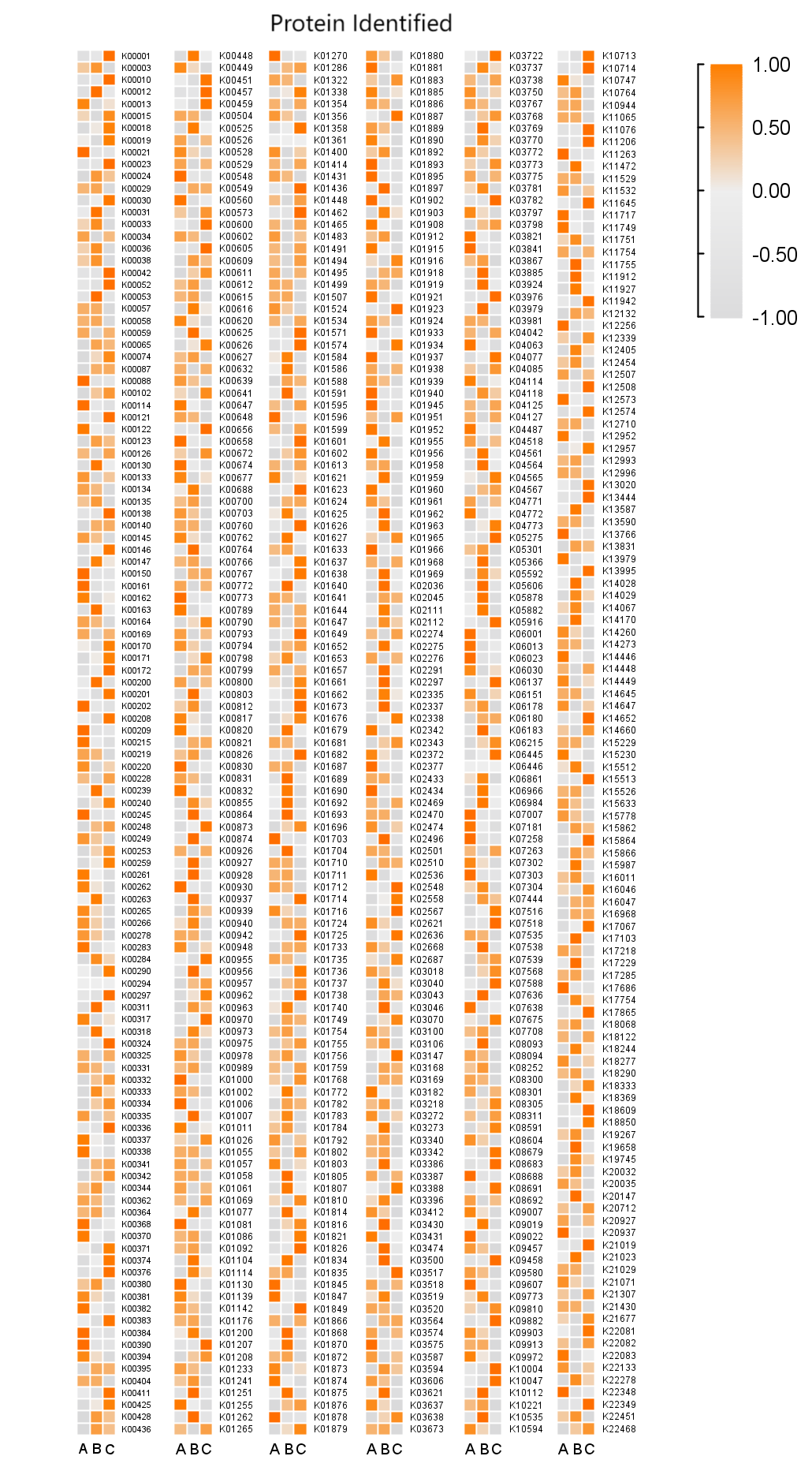

**Figure S3** Protein expression under tested conditions, Protein names in KEGG Orthology IDs

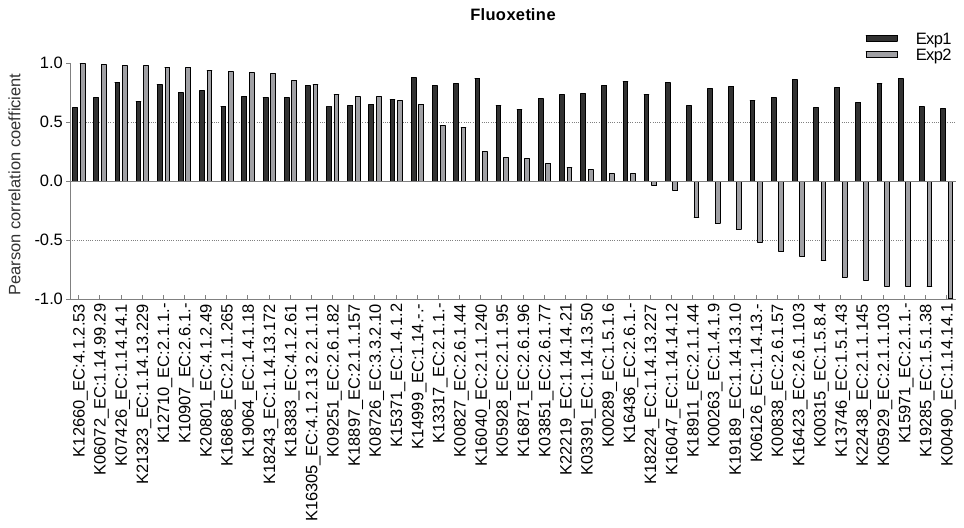

**Figure S4** Pearson correlation coefficients for fluoxetine with its corresponding ECs from both exp1 and exp2

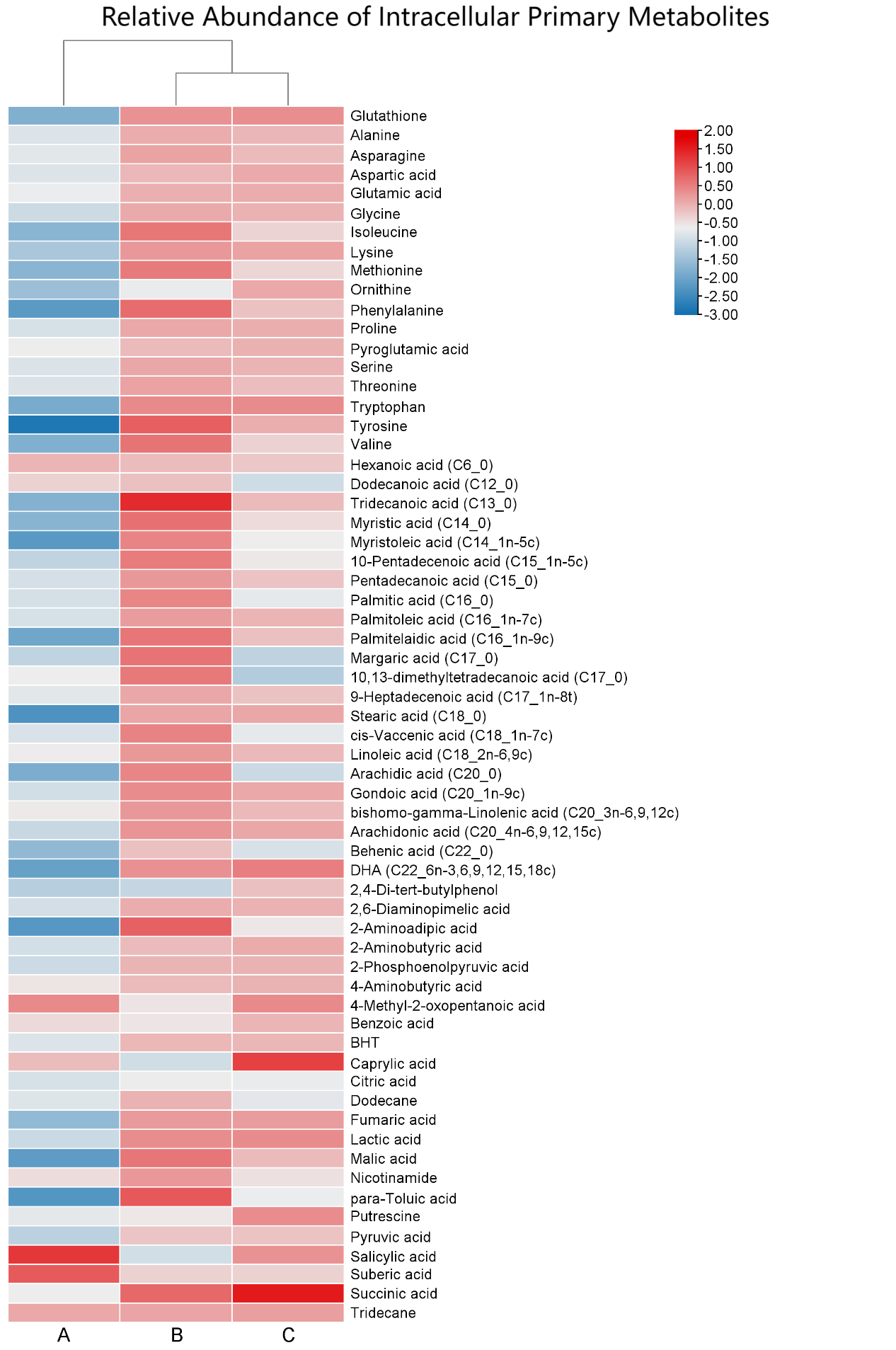

**Figure S5** The relative abundance of intracellular primary metabolites (amino acids, fatty acids, and organic acids) of microorganisms after 48 hours under conditions A, B, and C.
